# ssDNA hand-off from RPA to Polθ drives microhomology-mediated end-joining

**DOI:** 10.64898/2026.10.26.762034

**Authors:** Martha J. Longley, Hyunjee Lee, Owen Eby, Bhasker Dharavath, Yufan Wu, Abhilash Jayaraj, Felix Steinruecke, Haribabu Arthanari, Ralph Scully, Lucas Farnung, Joseph J. Loparo, Johannes C. Walter

## Abstract

DNA polymerase theta-mediated end joining (TMEJ) is the primary mechanism of double strand break repair in mitosis and becomes essential in BRCA-deficient cells. RPA, the ssDNA-binding protein that coats resected DNA ends, has long been thought to inhibit this pathway by blocking the initial microhomology search in TMEJ. Here we find instead that RPA plays an essential activating role in TMEJ. Structural, biochemical, and genetic analyses support a model in which Polθ binds directly to RPA at resected DNA ends and then competes with RPA for ssDNA. This releases ssDNA into Polθ’s helicase domain. We propose that threading two DNA ends through the Polθ dimer in this manner initiates the search for microhomology. Together, our findings establish a detailed mechanism of how proteins can harness RPA to access ssDNA for critical genome maintenance transactions.

## INTRODUCTION

DNA double-strand break (DSB) repair is essential for cell viability. A single unrepaired DSB is sufficient to trigger apoptosis, and incorrect DSB repair can lead to genomic instability and cancer. Polymerase theta-mediated end-joining (TMEJ) is a mutagenic repair pathway that is upregulated in many cancers (*1*). In this pathway, resected DSB ends are joined by Polθ using as few as two base pairs of microhomology, often leading to insertions, deletions and chromosomal translocations (*2*). Polθ consists of an N-terminal helicase domain (Polθ-H) that mediates Polθ dimerization, and a C-terminal error-prone DNA polymerase domain that is capable of bypassing lesions (Fig. S1A) (*3, 4*). These domains are joined by a large central region that is predicted to be disordered. Similar to homologous recombination (HR), Polθ acts on 3′ single-stranded DNA (ssDNA) overhangs generated by the MRE11-RAD50-NBS1 (MRN) complex and CtIP (*5-7*). The proteins RHINO and TopBP1 have been shown to promote its recruitment to these ends in mitosis, while its recruitment mechanism in interphase is less clear (*8, 9*). Polθ is thought to capture two ssDNA ends and search for microhomology via threading of ssDNA through the helicase domain (*10, 11*). FEN1, DNA polymerase delta (Polδ), and APEX2 have been implicated in cleavage of any 3′ flaps at the annealed junction (*12, 13*). Following flap removal, the polymerase domain of Polθ extends the annealed primer, which is then transferred to the more processive Polδ (*14*). Finally, the two strands are ligated (*12-14*). However, how Polθ gains access to ssDNA ends coated by replication protein A (RPA) and initiates the search for microhomology remains unclear.

RPA is the major eukaryotic ssDNA binding protein that plays central roles in DNA replication and repair (*15*). RPA is a heterotrimer composed of subunits RPA1, RPA2, and RPA3. Collectively, the three subunits contain six DNA-binding domains (DBDs) composed of oligonucleotide binding (OB)-folds, four of which (DBD-A, B, C, D) bind ssDNA (Fig. S1B). While RPA’s multiple DBDs mediate sub-nanomolar avidity of RPA for ssDNA, individual DBDs bind ssDNA with modest affinity, leading them to dynamically cycle on and off ssDNA, which is thought to allow for RPA displacement by other proteins (*15*). Due to RPA’s high abundance and high avidity for ssDNA, it affects virtually all genome maintenance processes that involve ssDNA intermediates. Some processes, such as nucleotide excision repair, single-stranded DNA annealing, and Okazaki fragment synthesis, are stimulated by RPA, while others, such as yeast MMEJ, appear to be inhibited by RPA (*16*). Accordingly, later work showed that partial knockdown of RPA stimulated TMEJ in mammalian cells (*5, 17, 18*). Consistent with this, Polθ’s helicase domain (Polθ-H) is able to remove RPA from ssDNA, which is thought to overcome RPA’s inhibitory effect on strand annealing (*5, 18, 19*). However, given that RPA is essential and highly abundant, assessing whether it promotes or inhibits a given process in cells is challenging.

To revisit the function of RPA in TMEJ, we depleted the protein from *Xenopus* egg extracts, a cell-free system that allows complete removal of essential factors. Surprisingly, we found that RPA is required for TMEJ in this system. Further analysis using structural, biochemical, and genetic approaches shows that Polθ’s helicase domain binds two RPA domains, DBD-C and DBD-D, in a manner that transfers ssDNA from RPA into the helicase pore. Thus, rather than simply counteracting RPA, Polθ uses RPA to both locate and capture ssDNA ends, setting the stage for repair.

## RESULTS

### RPA binding to Polθ is required for TMEJ

RPA is inherently difficult to study because it is essential. We therefore explored RPA’s role in TMEJ using nucleoplasmic *Xenopus* egg extracts (NPE), which support efficient DNA replication, most forms of DNA repair, and various signaling pathways in a quasi-cellular milieu (*20-24*). To monitor TMEJ, a linear DNA substrate containing a hairpin at one end and a ssDNA overhang with terminal microhomology at the other end was added to NPE (Fig. 1A). This substrate undergoes efficient dimerization (Fig. 1B, lanes 1-4), which requires both Polθ’s ATPase and DNA polymerase domains and is stimulated by Cyclin B-CDK1 (*25*). To assess how RPA affects joining in this system, we depleted it from NPE (Fig. 1C). Surprisingly, given its proposed inhibitory role in TMEJ, RPA-depleted extract was inactive for TMEJ, and this defect was rescued by the re-addition of recombinant *Xenopus* RPA expressed in bacteria (Fig. 1B, lanes 5-12, Fig. 1D). This finding suggests that RPA is essential for TMEJ in a vertebrate cell-free system.

**Figure 1.**
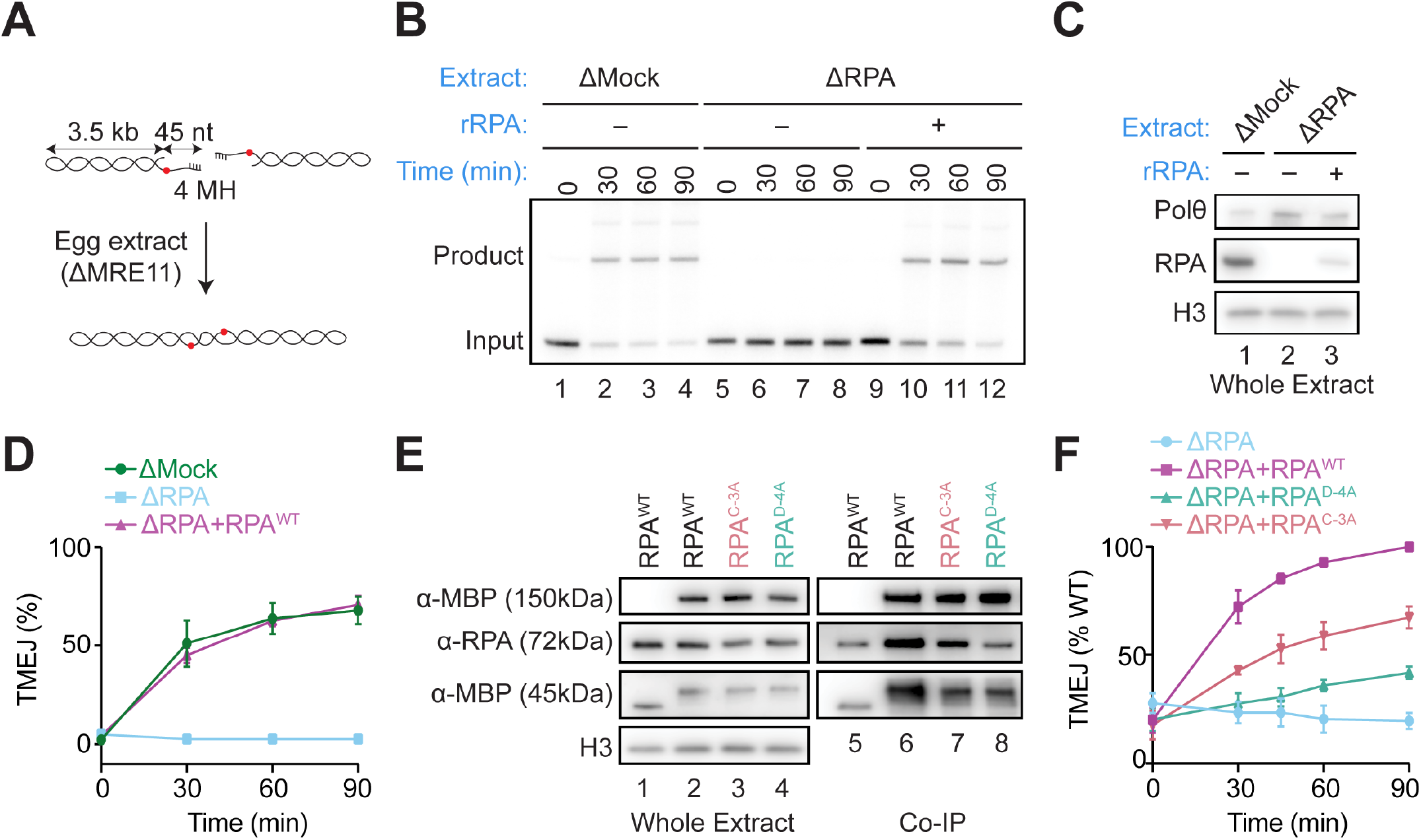
RPA binding to Polθ is required for TMEJ. **(A)** Diagram of the TMEJ assay in *Xenopus* Egg Extracts. Red dot = radiolabel. **(B)** Extracts were depleted of MRE11 to prevent excessive degradation of the substrate and supplemented with Cyclin B/CDK1 to simulate mitosis. Where indicated, extracts were depleted of RPA and reconstituted with recombinant *Xenopus laevis* RPA (rRPA). TMEJ was visualized using native agarose gel electrophoresis followed by autoradiography. Because RPA partially co-depletes Polθ, tobacco-expressed xPolθ was added to RPA-depleted extracts to match Polθ levels in mock-depleted extract. **(C)** Western blot of extracts used in (B). **(D)** Quantification of TMEJ repair products shown in (B). N=3 experiments. Mean±SEM. **(E)** Co-immunoprecipitation of RPA with Polθ-H. Tobacco expressed MBP-tagged xPolθ-H was immobilized on magnetic beads and added to RPA-depleted extracts supplemented with the indicated recombinant xRPA constructs. Beads were recovered, washed, and associated proteins were eluted and visualized using Western blotting. From left to right, whole extract and co-immunoprecipitation. (N=3). Lanes 1 and 5, MBP with no helicase domain controls. 150kD MBP blot shows full length MBP-xPolθ-H; in 45 kDa MBP blot, free MBP runs fast and an MBP-xPolθ-H proteolysis fragment migrates slower. **(F)** Quantification of TMEJ assay with doubly depleted extract (ΔMRE11 & ΔRPA) supplemented with the indicated RPA constructs, normalized to the TMEJ activity of wildtype RPA at 90 minutes. N=3 experiments. Mean±SEM. Representative gel and western blot shown in Figure S3C,D.

To understand how RPA promotes TMEJ, we consulted predictomes.org, a database of human protein-protein interactions nominated by *in silico* screening with AlphaFold-Multimer (AF-M) (*26*). RPA2 was one of the most confident Polθ interactors among ~270 core genome maintenance proteins surveyed (https://predictomes.org/view/ddr/group/202). We confirmed that AF-M predicts similar complexes in *Xenopus* and human (Fig. S2A-D). In the predicted structure, the helicase domain of Polθ (Polθ-H) binds in the ssDNA binding cleft of the DBD-D domain of RPA2 where it would compete directly with ssDNA (Fig. S2B, D). To address whether Polθ-H interacts with other RPA subunits, we modeled it with RPA1 and with RPA3. While no confident interaction was detected with RPA3, AF-M predicted that Polθ-H also interacts with DBD-C of RPA1 in *Xenopus* and humans (Fig. S2E-H). In these predictions, Polθ-H binds two spatially distinct surfaces of DBD-C without obstructing DBD-C’s ssDNA-binding groove (Fig. S2F, H). Based on this structural modeling, we hypothesized that RPA’s positive role in TMEJ is mediated through direct binding to Polθ-H.

To test the functional importance of the predicted RPA-Polθ-H contacts, we first asked whether RPA binds Polθ-H in NPE. RPA-depleted extracts were reconstituted with WT RPA and different RPA mutants and then supplemented with immobilized MBP-tagged Polθ-H and benzonase to degrade any endogenous DNA. As shown in Fig. 1E, RPA^WT^ was recovered with MBP-Polθ-H from the extracts, consistent with an interaction between the two proteins (Lanes 5-6). To determine which interfaces contribute to this interaction, we mutated conserved residues on RPA that are predicted to form salt bridges or hydrogen bonds with residues in Polθ-H. These substitutions generated *Xenopus* RPA^C-3A^ (D574A, E575A, S576A) which targets one of the two surfaces on DBD-C that are predicted to bind Polθ-H (Fig. S3A and S2H) and RPA^D-4A^ (D112A, D114A, E119A, R136A) which targets the Polθ-H-binding surface on DBD-D (Fig. S3B and S2D). RPA^D-4A^ bound Polθ-H at background levels, whereas RPA^C-3A^ exhibited partially defective binding (Fig. 1E, lanes 8 and 7), possibly due to the second Polθ-H-interacting surface on DBD-C that remained intact in the latter mutant (Fig. S2F, H). Importantly, RPA^C-3A^ and RPA^D-4A^ showed ~2- and ~5-fold reductions in cell-free TMEJ compared to RPA^WT^ at the 60 minute time point, respectively (Fig. 1F, Fig. S3C-D), but neither affected DNA replication (Fig. S3E-G), arguing against non-specific inactivation. These data indicate that direct interactions between Polθ-H and both DBD-C and DBD-D are required for efficient TMEJ.

### RPA recruits Polθ to DNA ends

We hypothesized that the RPA-Polθ interaction serves to recruit Polθ to RPA-coated ssDNA ends. To test this idea, a 5′ biotinylated ssDNA substrate with a free 3′ end was immobilized on streptavidin magnetic beads, incubated with egg extract, and recovered. The Polθ association with this substrate was lost in RPA-depleted extracts, and recruitment was restored with addition of recombinant RPA^WT^ (Fig. 2A, lanes 4-6). Compared to RPA^WT^, RPA^D-4A^ showed a severe defect in Polθ ssDNA recruitment, while the effect of RPA^C-3A^ was variable but on average reduced by ~50% (Fig. 2B, S4A), consistent with their effects on Polθ binding and TMEJ (Fig. 1E-F). Therefore, the stable recruitment of Polθ to DNA ends in egg extracts appears to depend on specific interactions between RPA and Polθ.

**Figure 2.**
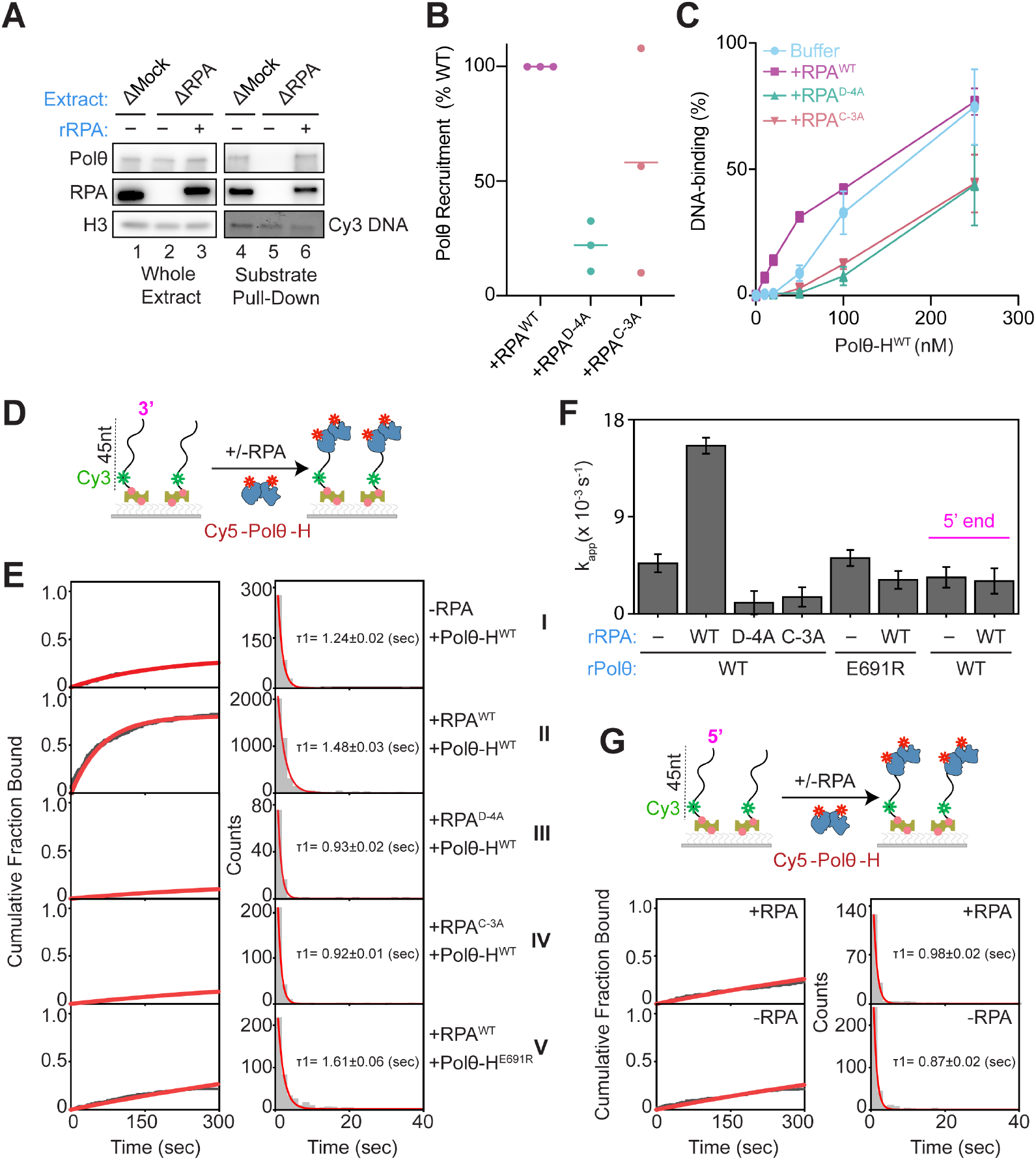
RPA recruits Polθ to DNA ends. (A)ssDNA pull-down assay: A 5′ biotinylated Cy3-labeled ssDNA with a free 3′ end was immobilized on magnetic beads and incubated with the indicated egg extracts and recombinant xRPA, as indicated. The substrate was washed and proteins were eluted and blotted. A representative western blot is shown of whole extract and proteins from recovered DNA. Cy3 is shown below as a loading control for ssDNA signal. N=3 experiments. (B) Quantification of pull-down assay with xRPA-depleted extracts supplemented with the indicated xRPA constructs, normalized to recruitment with xRPA^WT^. Mean of N=3 experiments. **(C)** Quantification of gel shift assay. Duplex DNA with a 3’ ssDNA overhang was incubated with hPolθ-H with and without hRPA constructs. N=3 experiments. Mean±SEM. Representative gel shown in Fig. S4B. **(D)** Diagram of Polθ-H single-molecule colocalization assay. 45 nucleotide 5′ biotinylated ssDNA was tethered to a coverslip and hRPA or buffer was flowed through. Next, Cy5-labeled hPolθ-H was flowed through and Cy5 colocalization was tracked. **(E)** Cumulative fraction of colocalized molecules (left) and dwell time histograms (right) for the indicated hPolθ-H and hRPA constructs. **(F)** Apparent first-binding rate constants (k_app_) for hPolθ-H under the indicated conditions, obtained by fitting the cumulative fraction of DNA molecules with a detected hPolθ-H binding event over time to a single-exponential association model. Bars indicate fitted (k_app_) values, and error bars indicate bootstrap standard errors. **(G)** Same assay as in (D) but with a 3′ biotinylated DNA (top) and cumulative fraction of colocalized hPolθ-H molecules (bottom left) and dwell times (bottom right) with and without hRPA.

We next asked whether the RPA-dependent recruitment of Polθ-H to ssDNA could be reconstituted with purified components. For these and subsequent biochemical experiments, we used human RPA and Polθ helicase. We measured human Polθ-H binding to a Cy3-labeled duplex DNA oligonucleotide with a 45 nucleotide 3′ ssDNA overhang using an electrophoretic mobility shift assay (Fig. S4B). We observed a marked increase in Polθ-H DNA binding in the presence of RPA^WT^, especially at low Polθ-H concentrations (Fig. 2C, purple vs. blue line). To produce human versions of RPA^C-3A^ and RPA^D-4A^, we mutated the analogous residues in human RPA (Fig. S3A-B, right panels). Binding of Polθ-H to the overhang substrate was reduced by human RPA^C-3A^ and RPA^D-4A^ to levels below what is seen in the absence of RPA (Fig. 2C, pink and green lines). Thus, RPA stimulates Polθ-H to bind single-stranded DNA, but only if the two proteins can interact.

To determine how RPA increases Polθ’s affinity for ssDNA, we used single-molecule imaging to quantify Polθ-H binding event frequency and dwell times. A Cy3-labeled ssDNA oligonucleotide was tethered to a coverslip at its 5′ end, and the colocalization of Cy5-labeled human Polθ-H was measured in the presence and absence of human RPA (Fig. 2D). Relative to the buffer condition, RPA increased Polθ-H binding events approximately 3-fold (Fig. 2E, I, II; Fig. 2F), indicating that RPA promotes Polθ-H association. In contrast, human RPA^D-4A^ and RPA^C-3A^ failed to increase binding event frequency or dwell time (Fig. 2E, III and IV, respectively; Fig. 2F), consistent with the decrease in shifted complexes in the EMSA (Fig. 2C). Instead, both mutants reduced the apparent first-binding rate of Polθ-H below that observed in the absence of RPA (Fig. 2F).

To further test the importance of the RPA-Polθ interface, we mutated Polθ-H residue E691, which is predicted to contact the DBD-D domain of RPA, to arginine (Fig. S3B, right panel). Polθ-H^E691R^ showed comparable ssDNA binding to Polθ-H^WT^ in the absence of RPA (Fig. S4D), but no increase in binding in the presence of RPA^WT^ in the single-molecule experiment or gel-shift assay (Fig. 2E, V, Fig. 2F, Fig. S5A-B). We attempted to purify Polθ-H constructs that disrupted its interaction with DBD-C but the constructs either inhibited binding to DNA (R460E) or destabilized the protein (K352A and K352S). Together, these data suggest that the stable association of Polθ-H with RPA-ssDNA is stimulated by the interaction between Polθ-H and RPA domains DBD-C and D. In the absence of these interactions, RPA inhibits Polθ-H from binding to ssDNA. Thus, by contacting RPA directly, Polθ-H converts RPA from an inhibitor of ssDNA binding into an enhancer of this process.

### Structure of the Polθ-H-RPA-ssDNA complex

To understand how Polθ-H leverages RPA to achieve efficient ssDNA binding, we determined a single-particle cryo-electron microscopy (cryo-EM) structure of the human Polθ-H-RPA-ssDNA complex. RPA, Polθ-H, and a 56-nucleotide ssDNA with six nucleotides of terminal microhomology were assembled and subjected to single-particle cryo-EM (Fig. S6A; Methods). Classification of the resulting dataset enabled a 3.4 Å resolution consensus reconstruction of a dimeric Polθ-H-RPA-ssDNA complex containing two copies of Polθ-H, two copies of the trimeric RPA core, and two stretches of 12-13 nucleotides of ssDNA (Fig. 3A-B, Fig. S6). Additional local refinements allowed the assembly of a composite map (Fig. S6A). Previously determined structures of Polθ-H with ssDNA and the RPA complex were placed into the composite reconstruction and adjusted locally (Fig. S7 and S8, Methods).

**Figure 3.**
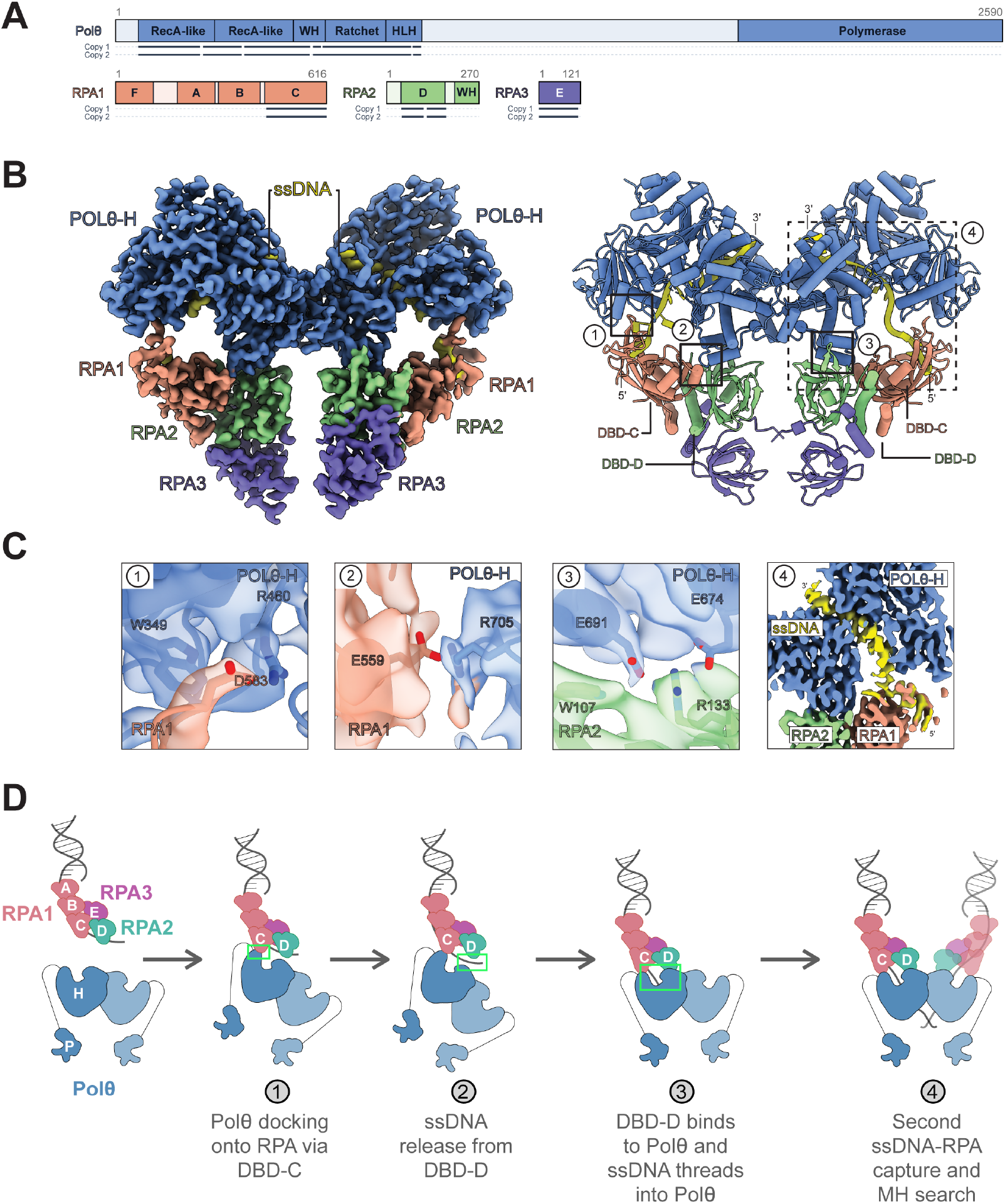
Cryo-EM structure of the dimeric Polθ-H-RPA-ssDNA complex. **(A)** Domain organization of human DNA polymerase θ, RPA1, RPA2, and RPA3. Modelled residues are indicated as black lines. **(B)** DeepEMhancer-sharpened composite map (Map G, left) and atomic model (right) of the Polθ-H-RPA dimer on microhomology-annealed ssDNA. **(C)** Atomic model and Coulomb potential map (map A) of the Polθ-H and RPA interfaces and path of ssDNA. 1, interaction of the Polθ-H RecA-like domain 2 with RPA1 DBD-C. 2, interaction of the Polθ-H ratchet domain with RPA1 DBD-C. 3, interaction of the Polθ-H ratchet domain with RPA2 DBD-D. 4, continuous path of ssDNA from Polθ-H core to the RPA1 DBD-C. **(D)** Model for how ssDNA is transferred from RPA to Polθ-H through contacts with DBD-C and DBD-D. DBD-F of RPA1 is omitted for clarity. The green box highlights events specific to each step.

In our structure, Polθ-H assembles into a tightly packed dimer with two-fold symmetry (Fig. 3B) that closely resembles a previously solved structure of Polθ-H on microhomology-annealed ssDNA (PDB: 9BHA, Cα RMSD 0.71Å across 766 pairs) (*11*). Each Polθ-H protomer binds one RPA trimerization core, positioning the two RPA molecules on the same rotational axis as the Polθ-H dimer (Fig. 3B). The structure confirms that Polθ-H contacts DBD-C through two spatially distinct interfaces. At one interface, disrupted in RPA^C-3A^, an acidic loop extending from RPA1 DBD-C forms electrostatic interactions with a basic groove on the surface of the Polθ-H RecA-like domain (Fig. 3B-C, view 1). Although not all the residues mutated in RPA^C-3A^ (Fig. S3A) are visible in the structure, D583 is clearly interfacial (Fig. 3C) and E584 is visible at high contour (Fig. S8G). In the second interface, a basic loop from the Polθ-H ratchet domain packs against an acidic helix on RPA1 DBD-C (Fig. 3B-C, view 2). At a third interface, the ssDNA binding groove of RPA2 DBD-D contacts two helices within the Polθ-H ratchet domain through a combination of aromatic and electrostatic interactions (Fig. 3C, view 3) that are disrupted in RPA^D-4A^ (Fig. S3B). We could only discern one of the residues (R133) mutated in RPA^D-4A^, as the disordered loop between V108 and V119 is not visible in our structure. The RPA-Polθ-H interfaces were similar for both protomers of the dimer (Fig. 3C and S8D). As in other structures of RPA (*27*), the N-terminal portion of RPA1 including DBD-F, DBD-A, and DBD-B, as well as the C-terminal RPA2 winged-helix domain, were not visible at high-resolution, indicating flexibility relative to the Polθ-H-RPA interface. However, lower-resolution density resembling DBD-B was visible at higher contours and in 2-D classes (Fig. S8A-B), consistent with its predicted location adjoining DBD-C.

Importantly, in each half of the complex, continuous ssDNA density was visible that extends from DBD-C into Polθ-H. Specifically, 3-4 nucleotides of the 5′ portion of the resolved ssDNA binds the DBD-C ssDNA binding groove in the previously described pose (Fig. S8E), while the remaining 9 nucleotides of ssDNA are threaded into the Polθ-H channel and engage its RecA-like helicase lobes (Fig. 3C, view 4) (*28*). The microhomology junction is visible in the structure with low pass filtering (Fig. S8C). No ssDNA density was associated with DBD-D, consistent with Polθ-H occupying the DBD-D ssDNA-binding cleft and preventing ssDNA rebinding to this domain.

These observations suggest an ordered hand-off mechanism by which Polθ-H accesses RPA-coated ssDNA (Fig. 3D). Because Polθ-H and ssDNA can co-occupy DBD-C but not DBD-D, we propose that Polθ-H first contacts RPA through DBD-C (Fig. 3D, step 1). Consistent with this possibility, molecular dynamics simulations indicated there is sufficient conformational flexibility in Polθ-H and RPA to allow Polθ-H docking onto DBD-C even while DBD-D remains bound to ssDNA (Fig. S9). Transient dissociation of ssDNA from DBD-D could then permit Polθ-H engagement with DBD-D, stabilizing the unbound 3′ ssDNA and enabling its entry into the helicase channel (Fig. 3D, steps 2 and 3), which resides near the site of ssDNA release (Fig. 3B). We propose that analogous events occur at both halves of the Polθ-H dimer, positioning the two resected ends for ATP-dependent microhomology search (Fig. 3D, step 4).

### RPA and Polθ contacts specify 3′-end loading and stimulate annealing

A prediction of our hand-off model is that RPA should stabilize Polθ-H binding only when a free 3′ end is available for threading into the helicase pore. To test this idea, we repeated the single-molecule co-localization experiment with ssDNA tethered at its 3′ end, leaving only a 5′ end accessible (Fig. 2G, top). In the absence of RPA, Polθ-H binding kinetics and dwell time to this 5′ substrate were comparable to the 3′ substrate (Fig. 2E-I, Fig. 2G, Fig. 2F). However, unlike what we observed with the 3′ substrate, addition of RPA did not increase Polθ-H binding to the 5′ substrate (Fig. 2F, 2G, and S4E). These data indicate that RPA enhances Polθ-H engagement with ssDNA in a manner that requires an accessible 3′ terminus. This polarity is consistent with a model in which RPA-Polθ contacts promote threading of 3′ ssDNA into the helicase pore.

The hand-off model also predicts that Polθ-H’s interaction with RPA should promote strand annealing by Polθ (*18*). To test this idea, we co-incubated two strands with 45 nucleotides of complementarity, one of which was labeled with Cy3, at concentrations that do not undergo spontaneous annealing (Fig. 4A, Fig. 4B, lanes 1 and 6). Polθ-H promoted annealing of these strands in a concentration dependent manner (Fig. 4B, lanes 1-5; Fig. 4C), and RPA further stimulated annealing, especially at subsaturating Polθ-H concentrations (Fig. 4B, lanes 6-10; Fig. 4C). In contrast, RPA^C-3A^ or RPA^D-4A^ inhibited Polθ-H-mediated annealing (Fig. 4B, lanes 11-20; Fig. 4C). Similarly, Polθ-H^E691R^, which disrupts binding to DBD-D, showed comparable annealing to Polθ-H^WT^ in the absence of RPA, but it supported much less annealing than Polθ-H^WT^ in the presence of RPA (Fig. 4D and Fig. S10A). Therefore, as seen above for ssDNA binding, the interaction between RPA and Polθ converts RPA from an inhibitor to a stimulator of Polθ-H-mediated strand annealing.

**Figure 4.**
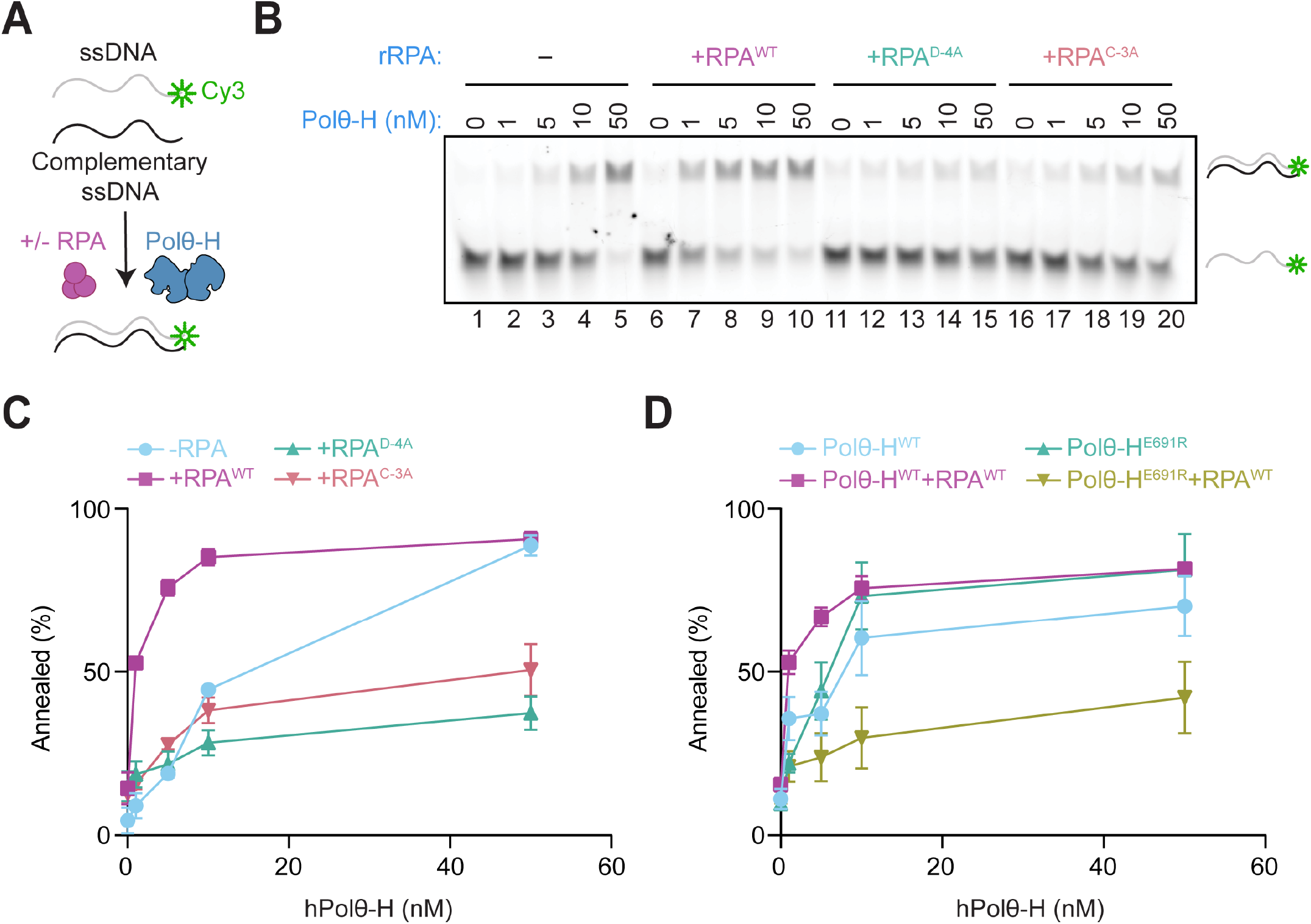
RPA promotes Polθ-mediated ssDNA annealing. **(A)** Scheme of annealing assay, where slower mobility of Cy3-labeled strand indicates annealing. **(B)** Representative gel of annealing in the presence of increasing concentrations of hPolθ-H and the indicated hRPA constructs. **(C)** Quantification of the experiment in (B). N=3, Mean±SEM. **(D)** Like C, with and without RPA and in the presence of Polθ-H^WT^ and Polθ-H^E691R^ as indicated. Representative annealing gel shown in Fig. S10A. N=3, Mean±SEM.

### Polθ-H strips RPA via interactions with the DBD-D and DBD-C domains

Polθ-H was previously shown to remove RPA from ssDNA in an ATP-dependent manner (*18, 19*). Our hand-off model predicts that this activity should be enhanced by the interactions between Polθ and RPA. To test this prediction, we tethered Cy3-labeled duplex DNA with a 70 nucleotide ssDNA overhang to a coverslip, prebound StayGold RPA (SgRPA), and then monitored SgRPA dissociation by loss of fluorescence intensity (Fig. 5A). Each ssDNA molecule was bound by ~2 RPA molecules (range 1-3; Fig. S10B-D). Upon addition of Polθ-H and ATP, the SgRPA dissociated from the DNA, which was observed as a step-wise decrease in intensity (Fig. 5B, compare panel II with panels I and III; Fig. S10C). In contrast, Polθ-H^WT^ failed to efficiently dissociate SgRPA^D-4A^ and SgRPA^C-3A^ (Fig. 5B, IV-VII), nor did Polθ-H^E691R^ remove SgRPA^WT^ (Fig. 5B, VIII). These data demonstrate that the interaction between the two proteins, as well as Polθ helicase activity, is essential for RPA stripping. Together, these experiments support our model that RPA actively promotes Polθ’s productive engagement with RPA-coated ssDNA, leading to ATP-dependent strand annealing and removal of RPA.

**Figure 5.**
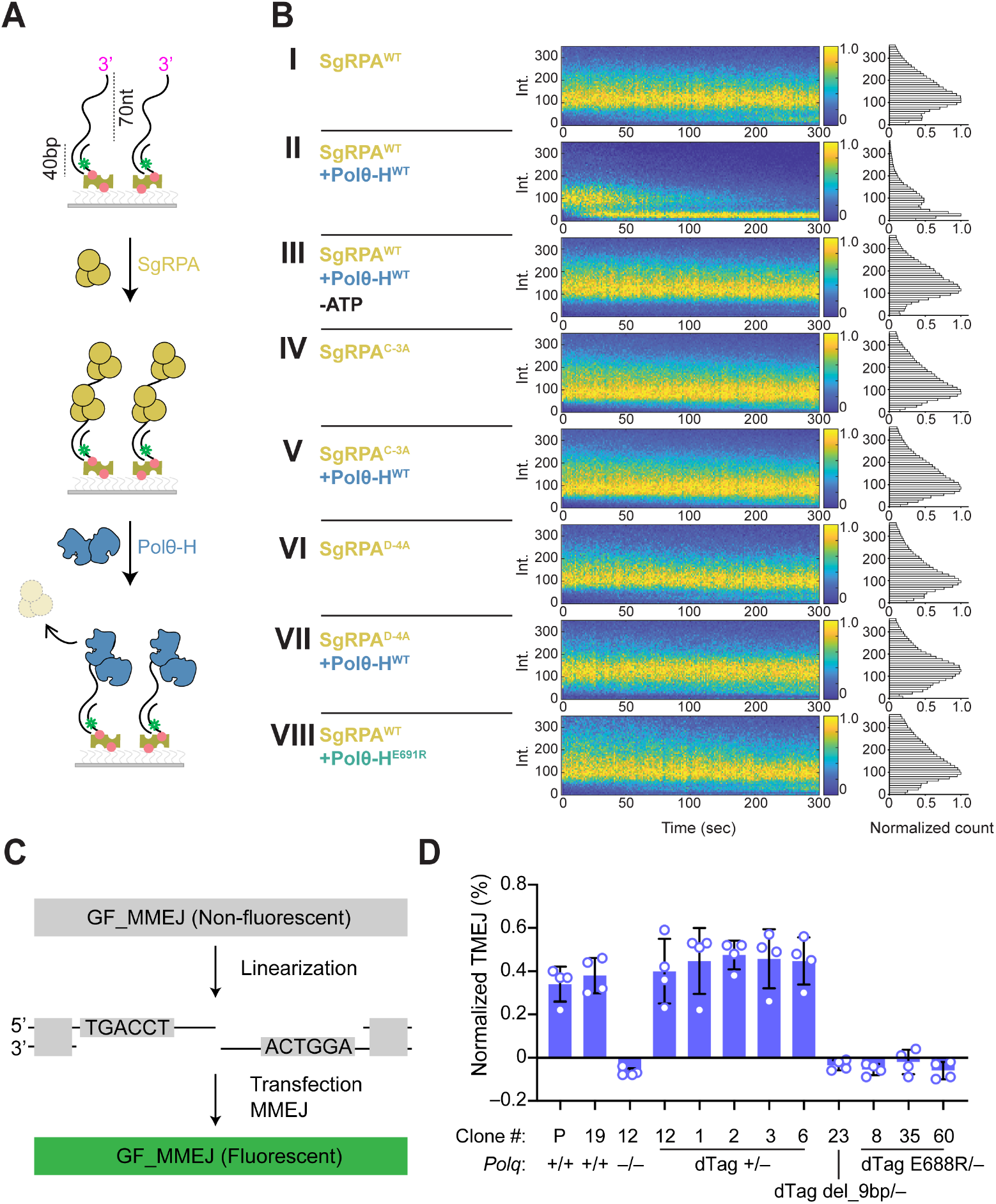
Polθ-RPA binding is necessary for RPA removal and TMEJ in cells. **(A)** RPA stripping assay. Labeled duplex DNA with a 3′ ssDNA overhang was incubated with StayGold WT or mutant RPA (SgRPA), and disappearance of SgRPA was subsequently tracked under the conditions indicated. **(B)** Time-resolved SgRPA fluorescence intensity histograms showing whether SgRPA dissociated under the indicated conditions based on the appearance of lower intensity signal. **(C)** Fluorescent reporter used to assay MMEJ in mES cells. **(D)** Normalized TMEJ measured using reporter in (C). See Methods for calculation of TMEJ and Supplemental Figure S11C for raw MMEJ values used to calculate TMEJ in these experiments. P: parental *Xrcc4*^fl/fl^ mES cell clone from which all *Polq* variants were derived. The last eight clones shown were all derived from founder clone *Polq*^dTag +/–^ clone #12. Mouse *Polq*^E688R^ corresponds to human *POLQ*^E691R^. *Polq*^dTag del_9bp/–^ clone contains a 9bp deletion within the *Polq* helicase-encoding region.

### The Polθ-RPA interaction is essential for TMEJ in cells

Finally, we asked whether the RPA–Polθ interaction is required for TMEJ in cells. Because RPA is essential and acts in many genome maintenance pathways, we mutated *Polq* rather than RPA. Using CRISPR/Cas9 gene disruption in mouse embryonic stem (mES) cells (*29*), we generated *Polq*^+/+^, *Polq*^−/−^ and *Polq*^+/−^ lines, and then targeted an in-frame C-terminal dTag-SMASh degron and HA epitope to the remaining allele of the hemizygous line (to generate *Polq*^dTag/−^). Immunoblotting for HA detected full-length Polθ at ~280 kDa in these cells but not in untagged parental cells (Fig. S11A). A second round of targeting introduced the helicase mutation E688R (equivalent to human E691R and *Xenopus* E821R). Three independent *Polq*^dTag E688R/−^ clones expressed full-length Polθ, as did four isogenic clones that had retained wild-type *Polq*; one further clone that carried a 9-bp deletion at the CRISPR/Cas9 target site expressed no detectable Polθ (*Polq*^dTag del_9bp/−^)(Fig. S11B). To measure TMEJ, we transfected a linearized reporter that restores *GFP* expression upon successful MMEJ (Fig. 5C) (*30*). We then scored Polθ-dependent TMEJ as the GFP^+^ cell frequency that was suppressed by the Polθ inhibitor ART558. Parental and *Polq*^dTag/−^ cells showed strong ART558-sensitive repair, whereas *Polq*^−/−^ and *Polq*^dTag del_9bp/−^ cells showed none, confirming the specificity of the assay (Fig. 5D, Fig. S11C). All three *Polq*^dTag E688R/−^ clones likewise showed no ART558-sensitive repair, whereas the four isogenic wild-type *Polq* clones were indistinguishable from parental cells. Thus, the interaction between the Polθ helicase domain and RPA DBD-D is essential for TMEJ in mammalian cells.

## DISCUSSION

RPA stimulates many genome maintenance processes that involve single-stranded DNA, including nucleotide excision repair, single-strand annealing, telomere extension, and Okazaki fragment priming and processing. In contrast, RPA has been viewed as an inhibitor of TMEJ that blocks Polθ access to ssDNA ends (*16*). Here, we use cryo-EM, biochemical analysis in egg extracts, mammalian cell biology, and single-molecule imaging to show that RPA’s dominant effect is to stimulate TMEJ by recruiting Polθ and coordinating ssDNA transfer to initiate repair.

Our findings support a multi-step “hand-off” mechanism that explains how ssDNA is transferred from RPA to Polθ (Fig. 3D). We show that Polθ uses two distinct surfaces on its helicase domain (Polθ-H) to interact with two of RPA’s ssDNA binding domains. Structural analysis shows that Polθ-H and ssDNA can co-occupy DBD-C, consistent with DBD-C serving as the site where Polθ-H first docks onto the RPA-ssDNA complex. Accordingly, MD simulations suggest that Polθ-H can bind DBD-C without clashing with ssDNA on DBD-D. Once bound to DBD-C, Polθ-H is poised to engage the DNA binding groove of DBD-D when the latter transiently releases the 3′ ssDNA end. This model is consistent with previous work showing that DBD-D binds dynamically to ssDNA (*31*). Once Polθ-H has engaged with DBD-D, the released 3′ ssDNA is captured by the helicase channel, which is located nearby. Finally, ATP-dependent Polθ translocation promotes the microhomology search by removing RPA, which we propose also renders ssDNA capture irreversible. The fact that RPA removal requires not only ATP hydrolysis but also the Polθ–RPA interaction argues against the prevailing view that RPA is stripped by the force of helicase translocation alone (*18, 19*). Overall, this ordered hand-off pathway represents a mechanism whereby a low abundance protein such as Polθ, which initially binds ssDNA with low affinity, can locate and capture ssDNA ends while simultaneously displacing RPA.

How do we reconcile our findings with prior observations that partial RPA depletion increases TMEJ in cells (*5, 18*)? Because RPA and Polθ can interact in the absence of DNA (Fig. 1E), one possibility is that the high level of RPA in nuclei sequesters a fraction of Polθ away from DNA ends. Partial RPA depletion could therefore increase the free pool of Polθ, boosting TMEJ, whereas when RPA is depleted fully, as in our extracts, TMEJ fails. Alternatively, reduced RPA levels may alter DNA damage signaling in a manner that promotes TMEJ. Either way, our finding that disrupting the RPA-Polθ interface abolishes TMEJ in cells strongly supports the conclusion that Polθ evolved to cooperate with – rather than simply oppose – RPA.

The concerted hand-off mechanism we propose requires Polθ to bind the terminal RPA at a DNA break. Therefore, the efficiency of hand-off likely depends on the extent of DNA end resection and thus the number of RPA molecules coating the terminal ssDNA. In mitosis, where TMEJ is a major DSB repair pathway, DNA ends are processed by MRN, but long-range resection is inhibited (*8, 32*). The resulting, short ssDNA overhangs are predicted to bind only one to two RPA molecules, greatly increasing the probability of a productive encounter between RPA and Polθ. In replication-coupled TMEJ, which was recently reported to depend on RPA, the amount of terminal ssDNA exposed is envisioned to be limited, which also creates an ideal substrate for RPA-dependent hand-off (*33*). Conversely, two-ended DSBs formed in S phase can undergo resection over several kilobase pairs of DNA, so most encounters between RPA and Polθ would be unproductive. However, such lesions are normally repaired by homologous recombination. We propose that in HR-deficient cells, which rely more heavily on TMEJ, Polθ transiently samples ssDNA until it finds the terminal RPA. In summary, the hand-off mechanism is tailored to Polθ’s physiological substrates while also allowing flexible repair of non-optimal substrates.

Several other genome maintenance proteins may engage in a similar hand-off mechanism with RPA. For example, RPA stimulates single-strand annealing by binding to RAD52 (*27, 34*), and RPA promotes the processing of 5′ flaps on Okazaki fragments by interacting with DNA2 (*35*). However, direct evidence that RAD52 and DNA2 release ssDNA from RPA by competition with RPA’s DBDs is lacking. In another example, the recombination mediator DSS1 binds RPA’s DBD-F domain and then competes with RPA’s ssDNA binding domains to facilitate BRCA2-dependent exchange of RPA for RAD51 (*36*). However, in this case, RPA appears to function purely as a barrier and not as a stimulatory platform. Finally, HELQ, a Polθ paralog that binds to and is stimulated by RPA(*37-39*), is predicted by AF-M to bind DBD-C and DBD-D in the same configuration as Polθ (*26*), strongly suggesting active hand-off of ssDNA from RPA. More work will be required to fully assess how closely different genome maintenance mechanisms mirror the Polθ-RPA interplay and whether RPA might sometimes stimulate processes it is generally considered to inhibit.

Our work also extends the concept that RPA leverages its asymmetric structure to support specific functions of its interacting proteins. RPA’s four primary DNA-binding domains (DBD-A, B, C, and D) are arrayed on ssDNA in the 5′ to 3′ direction (*35*). Consistent with this logic, Polθ’s specific interaction with DBD-C and DBD-D promotes engagement with 3′ ssDNA ends, a mechanism that also appears to be employed by telomerase (*40*). Thus, RPA not only facilitates client protein access to ssDNA, it also spatially constrains binding.

Polθ inhibitors have emerged as promising therapeutics in homologous-recombination deficient tumors (*41*). Based on our findings, targeting the RPA-Polθ interfaces could suppress Polθ function through a mechanism distinct from current inhibitors that block polymerase or ATPase functions. Such inhibitors could potentially be combined with existing Polθ inhibitors to suppress common resistance mechanisms.

## ACKNOWLEDGEMENTS

We thank members of the Loparo and Walter laboratories for discussions related to the manuscript. We thank Andrew Moreno and Brandon Case for help in overall single-molecule analysis and Ernst Schmid for discussions on AlphaFold analysis. We thank the Harvard Cryo-EM Center for Structural Biology for support with data collection.

## CONTRIBUTIONS

M.J.L. generated Figures 1, 2A,B, 3D, 4 and Supplementary Figures S3, S4A,B, S10A. H.L. generated Fig 2C-G, 5A,B; Supplementary Figures S4C,D, S5, S10C,D and Supplementary Table 1 and 2. O.E., F.S., and L.F. generated Figure 3, S1, S6, S7, S8 and Supplementary Table S3. B.D. and R.S. generated Figure 5D and Supplementary Figure S11. J.C.W. generated Figure S2. Y.W. contributed background data. A.J. and H.A. generated Supplementary Figure S9. M.L., H.L., J.C.W. and J.J.L. designed the study and wrote the manuscript with input from all authors. J.C.W. and J.J.L supervised the project.

## DATA AND CODE AVAILABILITY

Code for the initial single-molecule imaging was previously published (*42*). Cryo-EM maps have been deposited in the Electron Microscopy Data Bank (EMDB) under accession codes EMD-79697 (Map A, Polθ-H–RPA–ssDNA dimer consensus map), EMD-79704 (Map B, Polθ-H copy 1 focused refinement), EMD-79707 (Map C, Polθ-H copy 2 focused refinement), EMD-79716 (Map D, RPA copy 1 focused refinement), EMD-79717 (Map E, RPA copy 2 focused refinement) and EMD-79718 (Maps F and G, composite map and DeepEMhancer-sharpened composite map). The atomic model built into the composite map has been deposited in the Protein Data Bank (PDB) under accession code 39RH.

## METHODS

### AlphaFold2-Multimer Analysis

Structures were obtained from predictomes.org or modeled using Colabfold (*26, 43*) We then analyzed the relaxed structures using the ‘Protein interfaces, surfaces and assemblies’ (PISA), to identify which residues are predicted to mediate hydrogen bonds and salt bridges between RPA and Polθ (*44, 45*). Where noted, AlphaFold3 was used to model interactions involving phosphorylated proteins via the public webserver at https://alphafoldserver.com/ using default settings as exposed via the web interface (*46*).

### Expression and purification of recombinant proteins

#### *Xenopus* RPA

*Xenopus* RPA and associated mutants were purified as described previously, with the following modifications (*47*). Briefly, the plasmid [pRT019]pACE2-LacI-xRPA, which co-expresses *Xenopus* his-RPA1, RPA2, and RPA3 was transformed into E. coli C41(DE3) cells. 4L Luria broth supplemented with tetracycline was inoculated overnight without shaking at 37°C. The following afternoon, cultures were grown to an OD600 of 0.5-0.8 and induced with 0.3 mM IPTG. Cultures were subsequently shaken at 16°C for 16 hours. Cells were harvested and resuspended in lysis buffer (500 mM NaCl, 20 mM HEPES pH 7.5, 10 mM imidazole, 10% glycerol) and stored at −80°C. Pellets were thawed and resuspended with EDTA-free protease inhibitor cocktail (Roche) and lysed by sonication. Lysates were clarified by centrifugation (35,000 RPM Beckman Coulter Ti-45 rotor for 40 min at 4°C) followed by filtration through a 0.2-µm membrane. Clarified lysates were incubated with NiNTA resin pre-equilibrated in Lysis Buffer for 1h at 4°C while rotating. The resin was washed with 2×10 mL Lysis Buffer and eluted with 6×1mL Elution Buffer (500 mM NaCl, 20 mM Hepes pH 7.5, 400 mM imidazole, 10% glycerol). Peak fractions were pooled and diluted 10-fold into Buffer A (20 mM HEPES 7.5, 1 mM DTT, 0.5 mM EDTA, 10% glycerol). They were then centrifuged at 15,000×g for 5 min at 4°C. Next, they were loaded onto a 1 mL heparin column pre-equilibrated in 95% Buffer A and 5% Buffer B (20 mM HEPES 7.5, 1 mM DTT, 0.5 mM EDTA, 10% glycerol, 1 M KCl) and washed. RPA was eluted over a 20 mL gradient from 5% Buffer B to 100% Buffer B (RPA eluted around 300 mM KCl). Final fractions were pooled and concentrated using an Amicon Ultra centrifugal filter (30-kDa MWCO).

#### *Human* RPA

The plasmid p11d-tRPA (co-expressing human RPA1, RPA2, and RPA3) was transformed into *E. coli* BL21(DE3) cells and expressed in 6 L of 2×YT medium. Cultures were grown to an OD600 of 0.8 and induced with 1 mM IPTG, followed by incubation with shaking for 14–16 h at 16°C. Cells were harvested and resuspended in lysis buffer (25 mM HEPES pH 7.5, 100 mM NaCl, 0.5 mM EDTA, 1 mM DTT, 10% glycerol) supplemented with EDTA-free protease inhibitor cocktail (Roche), and lysed by sonication. Lysates were clarified by centrifugation (47,000 × g, 40 min) and the supernatant was filtered through a 0.45-µm membrane. The clarified lysate was loaded onto a HiTrap Blue HP column (Cytiva) pre-equilibrated in Lysis Buffer, washed with increasing KCl concentrations (up to 800 mM), and eluted with a linear NaSCN gradient (RPA typically eluted at ~750 mM NaSCN). Peak fractions were pooled and dialyzed overnight at 4°C against lysis buffer. The dialyzed sample was applied to a HiTrap Q HP column (Cytiva) and eluted with a linear NaCl gradient (100 mM–1 M). Fractions containing the RPA complex (RPA1/2/3) were pooled and concentrated using an Amicon Ultra centrifugal filter (30-kDa MWCO), followed by size-exclusion chromatography on a Superdex 200 Increase column (GE Healthcare). Purified RPA was analyzed by SDS–PAGE and stored at −80°C in storage buffer (25 mM HEPES, pH 7.5, 400 mM NaCl, 1 mM DTT, 10% glycerol, 0.5 mM EDTA). StayGold-RPA and mutant RPA variants were purified using the same protocol.

#### *Human* Polθ-H

The pE-SUMOstar vector (LifeSensors) encoding Polθ-H (aa 1–894) was transformed into E. coli Rosetta2(DE3)(pLysS) cells. Single colonies were used to inoculate LB and grown overnight at 37°C. The overnight culture was used to inoculate 6 L of autoinduction medium (1× Terrific Broth; USB Corporation) supplemented with 0.5% (w/v) glycerol, 0.05% (w/v) dextrose, 0.2% (w/v) α-lactose, and 10 mM MgCl_2_, and cultures were incubated with shaking for 55–60 h at 20°C. Cells were harvested and resuspended in Buffer A (50 mM HEPES, pH 8.0, 500 mM NaCl, 10% (v/v) glycerol, 5 mM β-mercaptoethanol (βME), 0.005% (v/v) Igepal CA-630), then lysed by sonication on ice. Lysates were clarified by centrifugation (47,000 × g, 40 min) and loaded onto Ni-NTA agarose resin (Qiagen) pre-equilibrated with Buffer A. The resin was washed with wash buffer (50 mM HEPES, pH 8.0, 450 mM NaCl, 10% (v/v) glycerol, 40 mM imidazole, 5 mM βME, 0.005% (v/v) Igepal CA-630), and Polθ-H was eluted with Buffer B (50 mM HEPES, pH 8.0, 450 mM NaCl, 10% (v/v) glycerol, 500 mM imidazole, 5 mM βME, 0.005% (v/v) Igepal CA-630). Fractions containing Polθ-H were pooled, treated with Ulp1 protease to remove the SUMO tag, and dialyzed overnight at 4°C against Buffer C (50 mM HEPES, pH 8.0, 100 mM NaCl, 10% (v/v) glycerol, 1 mM DTT, 0.005% (v/v) Igepal CA-630). The dialyzed sample was applied to a HiTrap Heparin HP column and eluted with a linear NaCl gradient (100–1,000 mM). Peak fractions were pooled, concentrated, and further purified by size-exclusion chromatography on a Superdex 200 Increase 10/300 GL column (GE Healthcare) equilibrated in SEC buffer (20 mM HEPES, pH 8.0, 150 mM sodium acetate, 150 mM KCl, 5% (v/v) glycerol, 1 mM DTT, 0.01% (v/v) NP-40). Purified protein was aliquoted, flash-frozen, and stored at −80°C. The E691R Polθ-H variant was expressed and purified using the same procedure.

### Preparation of labeled proteins

An N-terminal ybbR tag was introduced into the Polθ-H construct by site-directed mutagenesis. The ybbR-tagged Polθ-H was expressed and purified as described for wild-type Polθ-H, with the exception that buffer exchange and fluorescent labeling were performed prior to heparin chromatography. After Ni-NTA affinity purification, the eluate was exchanged into labeling buffer (20 mM HEPES, pH 8.0, 300 mM NaCl, 10 mM MgCl_2_, 10% glycerol, 1 mM DTT) using a PD-10 desalting column and concentrated. Site-specific labeling was carried out by incubating the protein (100 µM) with Sfp synthase (5 µM) and CoA-Cy5 (150 µM) overnight at 4°C in the dark. Ulp1 protease was included in the same reaction to cleave the SUMO tag. Excess free dye and enzymes were removed by heparin chromatography followed by size-exclusion chromatography, using the same conditions as for wild-type Polθ-H.

### *Xenopus* Egg Extract and in vitro assays

*Xenopus* egg extracts were prepared as described previously (*48*). Eggs were collected from adult female *Xenopus* laevis (Nasco Cat#4280). Center for Animal Resources and Comparative Medicine at Harvard Medical School (AAALAC accredited) cared for the frogs. All work performed for this study was in compliance with the rules and regulations determined by AAALAC and approved by the Institutional Animal Care and Use

### Replication assay

To license plasmids, high-speed supernatant (HSS) of total egg lysate was incubated at a concentration of 15 ng pBluescript per uL of HSS for 30 minutes at room temperature. Replication was initiated upon the addition of two volumes Nucleoplasmic Extract (NPE) with 1.93 mM DTT, 1.8mM ATP 18 mM phosphocreatine and 4.5 ug/mL creatine phosphokinase. Each reaction was supplemented with 0.16 uCi/uL of [α-^32^P] dATP (Perkin Elmer BLU512H500UC).

### TMEJ Substrate Preparation

Radiolabeled TMEJ substrate was prepared in the following manner. Pacesnapmre11, a derivative of pACEBac1, was linearized with BamHI and NotI (NEB). A 3.5 kb fragment was purified from a 1% agarose gel in TBE (89 mM Tris, 89 mM Boric acid, 2 mM EDTA pH 8.0). DNA was isolated from the gel using electroelution with 6-8kDa dialysis tubing (Fisher 08-670A) in TBE. DNA was precipitated (3 vol. 100% EtOH + 1/9 vol. 3 M sodium acetate pH 5.2) at −20°C for 1 hour or more. DNA was centrifuged at 15,000×g for 20 minutes and washed with 70% ethanol. Next, DNA was centrifuged at 15,000×g for 15 min. Ethanol was aspirated, the DNA was allowed to air dry for 10 minutes before resuspension in Milli-Q water.

The Hairpin Oligo (GGCCAGATGTCTTGCGTCCATGGACGCAAGACATCT) and Overhang Oligo (GATCAGTGAAGATCCTCACCTTCGGAGTACTCCTTCTGGTTAGAGC CGG) were phosphorylated using T4 PNK (NEB) for 30 min at 37°C. Both reactions were heat inactivated at 95°C for 5 minutes. The Hairpin Oligo was then annealed on ice in annealing buffer (1× T4 ligase buffer + 100 mM NaCl). The Overhang Oligo was also phosphorylated with [gamma-^32^P] (Perkin Elmer BLU502A250UC) by T4 PNK (NEB): 1 uL of 10 µM Overhang Oligo was radiolabeled with 3.8 µL 10 uCi/µL for 45 minutes at 37°C in PNK buffer. All reactions were heat killed at 95°C. The hairpin and hot and cold overhangs were ligated to the 3.5kb backbone using T4 Ligase for 6 hours at 16°C. Linear DNA was recovered using a Qiagen Gel Purification Kit (28706).

### TMEJ Assays

The ensemble gel-based end joining time course was performed similarly to a previous protocol (*49, 50*). For each reaction condition, the following were added to a final concentration in a mix of 70% NPE and 30% HSS: 2.16 mM ATP, 21.5 mM phosphocreatine, 5 ug/mL creatine phosphokinase and 30 ng/µL closed circular ‘carrier’ DNA. To initiate the joining reactions, the TMEJ substrate was added to reactions to a final concentration of 3 ng/µL. Reactions were carried out at room temperature. Unless otherwise indicated, extracts were supplemented with recombinant Cyclin B/CDK1 (Millipore Sigma) to a final concentration of 16 ng/µL and incubated for 10 minutes before initiating the reaction.

### Expression of proteins in tobacco protein expression system

Full length *Xenopus* Polθ was cloned into pALiCE01 and the sequence was confirmed using nanopore sequencing. Plasmids were purified from DH5α cells using ZymoPure II Plasmid Maxi Prep Kit (Zymogen). The proteins were expressed in the ALiCE Cell-Free Protein Synthesis (LenioBio) at a DNA concentration of 7 nM for 44 hours at 25°C, 700rpm. The samples were centrifuged at max speed at 4°C for 5 min following expression, and the supernatant was aliquoted, flash-frozen and kept at −80°C.

### Immunodepletions and rescue experiments

For immunodepletion of endogenous MRE11, we raised a rabbit antibody against *Xenopus* laevis MRE11. Recombinant xMre11 was overexpressed in E.coli, purified as inclusion bodies and injected into rabbits following standard procedures (Pocono Rabbit Farm). Immunodepletion using antibodies against *Xenopus* laevis Polθ (antigen: residues 1212 to 1506, Abgent) (*51*) and RPA (*52*) were performed as described previously. Briefly, 5 volumes of 1 mg/mL antibody or 2.5 volumes RPA-serum were incubated with 1 volume of Protein A Sepharose Fast Flow Beads (GE Healthcare) at 4°C overnight. Five volumes of pre-cleared egg extract were then mixed with one volume of the antibody-bound Sepharose beads. For Ku, Mre11, and Polθ depletion of NPE, three rounds of depletion were performed at 4°C for 1 hour each. For RPA, two rounds of 30 min and two rounds of 1h depletions were performed at 4°C. All were incubated on a rotating wheel.

In RPA experiments, recombinant RPA was added to a final concentration of 5 µM. In Polθ experiments, 1 volume of tobacco extract was added to 9 volumes of extract and incubated at room temperature for 10 minutes along with Cyclin B/CDK1 unless otherwise indicated. The DNA was subsequently added to initiate the reaction. In control lanes tobacco extract supplemented with an empty expression vector was added. Depletion efficiency was assessed by western blotting.

### Visualization and quantification of end joining and DNA replication Assay

At each timepoint indicated, 1 volume of reaction was quenched with 5 volumes replication stop buffer (80 mM Tris-HCl pH 8.0, 8 mM EDTA, 0.13% phosphoric acid, 10% Ficoll 400, 5% SDS, 0.2% bromophenol blue) with 20 µg of proteinase K (Roche 3115879001). Proteins were digested for one hour at 37°C. Samples were run on 0.8% agarose gels in TBE buffer (89 mM Tris, 89 mM Boric acid, 2 mM EDTA pH 8.0). Gels were dried atop a positive charged membrane (GE/Cytiva Hybond-XL or Roche 11417240001) to retain nucleic acids. Once dried, gels were exposed to phosphor screens. Screens were then imaged on the Typhoon FLA 700 PhosphorImager (GE Healthcare). DNA synthesis or joining was quantified using ImageJ. Background intensity was measured in a DNA-free lane using a region of interest of the same size and shape at the corresponding position and subtracted from each band intensity. Annealing (%) was calculated as the background-corrected intensity of the joined products divided by the sum of the background-corrected intensities of the joined products and monomeric substrate, multiplied by 100. Data represent the mean ± SEM from three independent experiments.

### SDS-PAGE and immunoblotting of samples from *Xenopus* egg extract experiments

Protein samples were boiled in Laemmli buffer (50 mM Tris-HCl pH 6.8, 2% SDS, 10% glycerol, 0.1% bromophenol blue, 5% β-mercaptoethanol). Unless noted, samples were run in 4-15% Mini-PROTEAN TGX Precast Protein Gels (Bio-Rad #4561086) using Tris-Glycine-SDS Running Buffer (25 mM Tris-HCl pH 8.3, 192 mM glycine, 0.1 % SDS). The samples were run alongside EZ-Run Prestained Rec Protein Ladder (Fisher BioReagents #BP36031) and the Color-coded Prestained Protein Marker, High Range (Cell Signaling Technology) to track the size of the protein bands. Protein purification gels were stained using InstantBlue stain (Novus ISB1L) for 1 hour at room temperature with gentle agitation. For western blotting, proteins were transferred to PVDF membranes (Thermo Scientific #88518) in Transfer Buffer (25 mM Tris pH 8.5, 192 mM glycine, 20% methanol). Membranes were subsequently blocked in PBST with 5% (w/v) nonfat milk powder (Bob’s Red Mill) for 30 minutes at room temperature with gentle shaking. Membranes were washed 3×5 minutes in PBST before incubation with primary antibodies diluted into PBST with 1% (w/v) BSA overnight at 4°C with gentle agitation. Membranes were washed with PBST for 3×5 minutes before incubating with Peroxidase IgG Fraction Monoclonal Mouse Anti-Rabbit IgG, Light Chain Specific (Jackson ImmunoResearch) diluted 1:5000 in PBST supplemented with 5% (w/v) nonfat milk for 1 hour at room temperature with agitation. Membranes were washed 3×5 minutes with PBST and developed using ProSignal Pico ECL Spray (Prometheus Protein Biology Products #20-300S) and imaged using an Amersham Imager 600 (GE Healthcare). The following antibody dilutions were used:

RPA: 1:7000 (*52*)

H3: 1:500, Cell Signaling #9715, RRID: AB_331563

Polθ: 1:500 (*51*)

### Pull-Down Assay

1. volume of streptavidin-coated magnetic beads (Dynabeads M-280, Invitrogen 11206D) was washed and incubated with 4 volumes Binding Buffer (10 mM Tris pH 7.4, 2 M NaCl, 1 mM EDTA) supplemented with 8.3 nM 5’ biotinylated ssDNA (/5Biosg/TCAGATATCTAGCTTAAAT/iCy3/GGTGGCTCGCCGGCACTGCACTACCCGGGTATTTTTTTTTTTTTTTTTTTTTTTTTTTTTTTTTTTT TTTT). Beads were rotated for 1 h at 4°C. The beads were washed 2 × with Binding Buffer and 1× in Blocking Buffer (250 mM sucrose, 10 mM HEPES-KOH pH 7.7, 50 mM KCl, 2.5 mM MgCl2, 0.02% Tween-20, 0.25 mg/mL BSA). Beads were resuspended in 1 volume Blocking Buffer and supplemented with 2 volumes extract. For each reaction condition, the following were added to a final concentration in extracts (70% NPE and 30% HSS): 2.16 mM ATP, 21.5 mM phosphocreatine, 5 ug/mL creatine phosphokinase, 16 ng/µL Cyclin B/CDK1 (Millipore Sigma). Where indicated, RPA was added back to a final concentration of 5 nM and Polθ expressed in tobacco extract was added to a final concentration of 10%. Samples were incubated for 30 min on a rotating wheel at room temperature. 1 µL whole extract was diluted into 19 µL Laemmli Buffer. At 30 min, the samples were diluted 1:5 into streptavidin beads washed and resuspended in Blocking Buffer. Samples were washed 3×200 µL cold Wash Buffer (20 mM HEPES-KOH pH 7.7, 100 mM KCl, 5 mM MgCl2, 0.03% Tween-20, 0.25 mg/mL BSA). To elute proteins 15 µL 1 × LB was added to the pellets and boiled for 2 minutes at 95°C. 3 µL whole extract and 6 µL pull-down was loaded for western blot.

### Immunoprecipitation Assays

A construct with MBP-tagged *Xenopus* laevis Polθ Helicase (residues 1-1021) and MBP alone were cloned into pALiCE01. Plasmids were purified from DH5α cells using ZymoPure II Plasmid Maxi Prep Kit (Zymogen). The proteins were expressed in the ALiCE Cell-Free Protein Synthesis system (LenioBio) at a DNA concentration of 5 nM for 44 hours at 25°C, 700 rpm. Per reaction, 10 µL ALiCE extract was incubated with 5 µL Anti-MBP Beads (NEB #8037S) pre-washed in IP Wash Buffer (10 mM HEPES-KOH pH 7.7, 50 mM KCl, 2.5 mM MgCl_2_, 250 mM sucrose, 0.1 mg/mL BSA, 0.05% Tween

20) for 1 hour at 4°C. Following incubation, beads were washed with 3×200 µL IP Wash Buffer and resuspended in 10 µL 50% RPA-depleted HSS supplemented with 5 nM indicated xRPA constructs or buffer. Reactions were supplemented with Benzonase (Sigma 70746-3) and MgCl2 (final concentration 1 mM) and incubated on a rotating wheel for 1 h at 4°C. A whole extract sample was diluted 1:29 in Laemmli Buffer. The remaining extracts were washed 3×200 µL Wash Buffer (10 mM HEPES-KOH pH 7.7, 500 mM KCl, 2.5 mM MgCl_2_, 0.05% Tween-20, 0.1 mg/mL BSA). Samples were eluted in 15 µL Laemmli Buffer and boiled at 95°C for 2 minutes. 3 µL whole extract and 6 µL immunoprecipitated extract were analyzed by western blot.

### Electrophoretic mobility shift assay

#### Preparation of DNA Substrates

Three DNA oligonucleotides were assembled and ligated to generate the EMSA substrate: a 5′-phosphorylated 60-nt oligonucleotide (5′-GTCTGGCCTGTCTTAAGTGA AGATCCTCACCTTCGGAGTACTCCTTCTGGTTAGAGCCGG-3′), an 85-nt Cy3-labeled oligonucleotide (5′-GAGCTCA/iCy3/AATGTCGGGCAATCAGGTGCGACAATCTAT CGATTGTATGGGAAGCCCGATGATGCATTGCATGAGATGCTTGCGT C-3′), and a 100-nt biotinylated oligonucleotide (5′-TAAGACAGGCCAGACGACGCAAGCATCTCA TGCAATGCATCATCGGGCTTCCCATACAATCGATAGATTGTCGCAC CTGATTGCCCGACATTATGAGCTC/3Bio/-3′). Oligonucleotides were combined at a 1:1:1 molar ratio in T50 buffer (10 mM Tris-HCl, pH 8.0, 50 mM NaCl), heated to 95°C for 2 min, and slowly cooled to 16°C to form a three-piece annealed complex. The annealed complex was ligated with T4 DNA ligase in 1× T4 DNA ligase reaction buffer at 16°C overnight. Ligation products were resolved on a 6% native polyacrylamide gel (TBE). Samples were supplemented with 50% glycerol and electrophoresed at 5 W for 2 h. The band corresponding to the expected ligation product was excised and DNA was recovered by gel extraction, followed by elution in TE buffer (10 mM Tris-HCl, pH 8.0, 0.1 mM EDTA).

### EMSA

Reactions were assembled in binding buffer (25 mM HEPES, pH 8.8, 30 mM NaCl, 1 mM DTT, 0.1 mg/mL BSA, 0.01% NP-40, 10% glycerol) containing 20 nM Cy3-labeled DNA substrate (prepared as described above). DNA was incubated in the absence or presence of 50 nM RPA (WT or the indicated variants) for 30 min at room temperature, followed by addition of Polθ-H and a further 30 min incubation. Reactions were mixed with SDS-free 6× loading dye (Gel Loading Dye, Purple; no SDS) and resolved on a 2% agarose gel in TBE. Electrophoresis was performed at 150 V for 50 min at 4°C. Gels were imaged by scanning Cy3 fluorescence on a Typhoon 5 scanner.

### Annealing Assay

Annealing assay was adapted from a previously published protocol (*39*). 15 µL reactions were assembled in Annealing Buffer (25 mM Tris-HCl pH 8.0, mM ATP, 2 mM MgCl2, 1 mM DTT, 50 mM NaCl, 0.1 mg/mL BSA). Oligo 1 (/5Cy3/AGCTACCATGCCTGCACGAATTAAGCAATTCGTAATCATGGTCATAGT) was added to a final concentration of 5 nM. RPA was diluted in Annealing Buffer and added to a final concentration of 40 nM. Polθ-H was diluted in Annealing Buffer and added at indicated concentrations and reactions were placed at 37°C. Reactions were initiated by the addition of Oligo 2 to a final concentration of 5 nM (AGCTATGACCATGATTACGAATTGCTTAATTCGTGCAGGCATGGTAGT). Reactions were stopped after 12 minutes with 6 µL Stop Solution (0.2% SDS, 30% glycerol, 150 mM EDTA, Orange G) supplemented with 100-fold excess of cold oligo 1 (AGCTACCATGCCTGCACGAATTAAGCAATTCGTAATCATGGTCATAGT). Proteins were digested with 20 µg proteinase K (Roche 3115879001) and incubated for 10 min at 37°C. Products were resolved on a 10% native polyacrylamide gel and imaged for Cy3 fluorescence on a Typhoon 5 imager. Band intensities were quantified using ImageJ. Background intensity was measured in a DNA-free lane using a region of interest of the same size and shape at the corresponding position and subtracted from each band intensity. Annealing (%) was calculated as the background-corrected intensity of the annealed product divided by the sum of the background-corrected intensities of the annealed product and unannealed ssDNA, multiplied by 100. Data represent the mean ± SEM from three independent experiments.

### Single-molecule Microscopy

Single-molecule imaging was carried out on a home-built objective-type TIRF platform (Olympus IX-71) following established configurations (*53-55*). Excitation was delivered with 488-, 532-, and 641-nm lasers depending on the fluorophore (StayGold, Cy3, and Cy5, respectively). Emission was separated into Cy3 and Cy5 channels and recorded on an EMCCD camera (Hamamatsu ImageEM 9100-13) at high EM gain. Channel-to-channel mapping was obtained using 0.1-µm multicolor fluorescent beads (TetraSpeck) and applied to dual-color movies. For StayGold/Cy3 experiments, the two signals were acquired in the Cy3 channel by alternating 488- and 532-nm illumination. Fields of view were selected using a motorized stage. Laser power at the sample plane was measured under TIRF alignment (Coherent FieldMate with OP-2 VIS) and reported as through-objective power unless stated otherwise.

### Preparation of Flow Cell

Flow cells were built as a single-channel microfluidic chamber using PEG-passivated coverslips containing a fraction of biotin-PEG and a pressure-sensitive spacer (SecureSeal, Grace Bio-Labs). A drilled glass slide served as the manifold for inlet/outlet connections, which were secured with epoxy. Solutions were introduced by withdrawing with a syringe from the outlet line. Prior to imaging, surfaces were functionalized with streptavidin (1 mg/mL in PBS), rinsed with PBS, and incubated with biotinylated DNA substrates (50 pM in PBS) to obtain sparse immobilization. Excess DNA was removed by washing with PBS.

### Single-molecule co-localization assay

Cy3-labeled 45-nt ssDNA (5′-/5BiosG/AGTGAAG/iCy3/ATCCTCACCTTCGGAGTACTCCTTCTGGTTAGAGCCGG-3′) was immobilized on PEG/biotin-PEG–functionalized coverslips in a microfluidic flow cell prepared as described above. After immobilization, the chamber was incubated with BSA (1 mg/mL) for 10 min to reduce nonspecific surface binding and then washed with PBS. Where indicated, RPA (WT or the indicated variants) was introduced at 50 nM, incubated for 5 min, and unbound RPA was removed by washing with PBS. Cy5-labeled Polθ-H (20 nM) was subsequently introduced in imaging buffer containing 25 mM HEPES (pH 8.8), 30 mM NaCl, 10% glycerol, 1 mM DTT, 0.1 mg/mL BSA, 0.01% NP-40, 8 mM PCA, 160 nM PCD, and 2 mM Trolox. Binding was monitored by colocalized Cy3 and Cy5 fluorescence signals. Images were acquired continuously at 5 frames/s with alternating excitation (one frame at 532 nm followed by four frames at 641 nm). For each experiment, a single field of view was recorded for 5 min. Laser powers were set to ~0.7 mW for both 532-nm and 641-nm excitation.

### Single-molecule RPA removal assay

A separate three-oligonucleotide assembly was used to prepare substrates for the single-molecule RPA removal assay: a 5′-phosphorylated 90-nt oligonucleotide (5′-/5′Phos/TCAGGTGCGACAATCTATCGAGAGAGAGGAGAAGAAGAAGAAGAAGAGAGAGGAGAAGAAGAAGAAGAGAAGAGAAGGAGGAGAGACCGG-3′), a 20-nt Cy3-labeled oligonucleotide (5′-GAGCTCA(iCy3)TAATGTCGGGCAA-3′), and a 40-nt biotinylated oligonucleotide (5′-CGATAGATTG TCGCACCTGA TTGCCCGACA TTATGAGCTC/3Bio/-3′) following the protocol described in EMSA substrate preparation. To visualize real-time dissociation of RPA from DNA, StayGold was fused to the C terminus of RPA1. StayGold-RPA was expressed and purified as described for recombinant human RPA. Single-molecule RPA removal assays were performed using 532-nm and 488-nm excitation for Cy3 and StayGold, respectively. DNA substrates were prepared in flow cells as described for the single-molecule co-localization assay. StayGold-RPA (2 nM) was introduced in imaging buffer containing 40 mM HEPES (pH 8.0), 50 mM NaCl, 5 mM MgCl_2_, 1 mM ATP, 1 mM DTT, 0.2 mg/mL BSA, 8 mM PCA, 160 nM PCD, and 2 mM Trolox. After a 5-min incubation, unbound StayGold-RPA was washed out with imaging buffer, and Polθ-H (50 nM) was subsequently introduced. For the ATP-free condition, ATP was removed after RPA loading by washing the flow cell with ATP-free imaging buffer before addition of Polθ-H. Polθ-H was subsequently introduced in ATP-free imaging buffer. StayGold-RPA removal was monitored as the loss of StayGold fluorescence at Cy3-marked DNA molecules. Images were acquired with a 200-ms exposure time using alternating excitation (one frame at 488 nm followed by one frame at 532 nm); Laser powers were set to ~0.7 mW for both 488-nm and 532-nm excitation.

### SINGLE-molecule photobleaching test

The single-molecule photobleaching assay was performed using the same flow-cell preparation and DNA/StayGold-RPA loading procedure as in the StayGold-RPA removal assay, unless otherwise noted. After binding of StayGold-RPA and removal of unbound protein, imaging was carried out in buffer lacking the oxygen-scavenging and anti-photobleaching components (PCA, PCD, and Trolox). Samples were imaged using the same alternating excitation scheme as in the removal assay (one frame at 488 nm followed by one frame at 532 nm), with the 488-nm laser power increased to 5 mW (532 nm unchanged) to achieve >90% photobleaching of StayGold-RPA. All other imaging settings were identical to those used for the removal assay.

### Analysis of single-molecule data

Single-molecule movies were processed in MATLAB using custom scripts and, where indicated, a published automated workflow (*42*). The Cy3 and Cy5 channels were aligned using an affine mapping obtained from multicolor fluorescent beads. Background-corrected fluorescence trajectories were extracted at DNA positions and normalized. Binding events were identified by change-point analysis of normalized Cy5 trajectories and classified using intensity, spot-size, and spatial criteria, with a maximum distance of 1.2 pixels from the corresponding Cy3-labeled DNA position. Dwell times were calculated from the timestamps marking the beginning and end of each detected bound interval. For association analysis, only the first detected binding event at each DNA molecule was counted. The cumulative fraction bound was calculated as the number of DNA molecules with at least one detected binding event before each time point divided by the total number of analyzed DNA molecules. These curves were fitted by nonlinear least squares in MATLAB to F(t) = F_max_[1 − exp(−k_app_ t)], where Fmax is the fitted plateau and k_app_ is the apparent first-binding rate constant. Dwell-time histograms were fitted in Origin using a single-exponential decay function (ExpDec1), y = y_0_ + A_1_ exp(−t/τ), where y_0_ is the baseline offset, A_1_ is the amplitude, and τ is the decay time constant. Reported uncertainties in τ represent the standard errors of the fitted parameter.

StayGold–RPA datasets were analyzed with the automated MATLAB pipeline for spot detection and extraction of background-corrected intensities (*42, 56*). Co-localization was called using assay-specific intensity and displacement criteria (maximum distance 1.2 pixels), requiring a minimum dwell of two frames and enforcing a two-frame separation between successive events. Thresholds were selected based on control ROIs lacking DNA signal to minimize nonspecific calls. Processed traces and event annotations were exported for downstream analyses. DNA molecules with detectable StayGold–RPA signal in the first analyzed frame were included, and their fluorescence intensities were followed throughout the analyzed interval, including after signal loss. Time-resolved intensity histograms were displayed as heatmaps, with bin counts at each time point divided by the maximum bin count at that time point. For the RPA removal assay, fluorescence intensity values were pooled over the analyzed time interval to generate intensity histograms in Origin. Histogram bin counts were rescaled to the range 0–1.

Photobleaching steps in selected StayGold–RPA fluorescence trajectories were identified using custom MATLAB scripts incorporating smoothing and change-point analysis. Downward intensity transitions, including the final transition to baseline, were counted to estimate the number of DNA-bound RPA molecules. For each independent experiment, the probability of each step count was calculated as the number of trajectories with that step count divided by the total number of analyzed trajectories. Data represent the mean ± SEM from four independent experiments.

### Mouse embryonic fibroblast and stem cell culture

Mouse embryonic fibroblast (MEF) feeder layers were prepared 1–2 days prior to embryonic stem (ES) cell plating. Irradiated MEFs were thawed, resuspended in complete MEF medium (DMEM, 15% fetal bovine serum (FBS), 0.8mM β-mercaptoethanol, 2mM L-glutamine, 100U/mL penicillin, 100µg/mL streptomycin), and seeded onto 6-well plates pre-coated with 0.2% gelatin. Feeder cells were allowed to form a confluent supportive layer before use. Conditional *Xrcc4* mouse ES cells, in which both *Xrcc4* alleles harbor loxP sites flanking exon 3 (floxed exon 3), were thawed and plated directly onto the prepared MEF feeder layers. Cells were maintained in ES cell medium consisting of Dulbecco’s Modified Eagle Medium (DMEM; with L-glutamine, 4.5 g/L glucose, and sodium pyruvate; Corning, Cat. No. 10-013-CV) supplemented with 15% FBS, 20 mM HEPES (pH 7.6), 0.1 mM β-mercaptoethanol, 500 U/mL recombinant leukemia inhibitory factor (rLIF), 1× MEM non-essential amino acids (NEAA), 2 mM L-glutamine, 100 U/mL penicillin, 100 µg/mL streptomycin, and 1 mM sodium pyruvate. Cells were cultured at 37°C in a humidified incubator with 6% CO_2_ and passaged every 2–3 days to prevent overconfluency and differentiation. All cell lines were routinely screened for mycoplasma contamination using the MycoAlert Mycoplasma Detection Kit (Lonza), and only mycoplasma-free cultures were used for experiments. All *Polq* hemizygous and mutant clones were derived from the parental conditional *Xrcc4* ES cell line and were cultured as described above.

### sgRNA synthesis and purification

Single guide RNAs (sgRNAs) were synthesized by in vitro transcription using the EnGen sgRNA Synthesis Kit (New England Biolabs, Cat. No. E3322), following the manufacturer’s protocol. Briefly, DNA templates containing the T7 promoter and target-specific sgRNA sequences were incubated with T7 RNA polymerase. After transcription, residual DNA templates were removed by DNase I treatment. sgRNAs were purified using the RNA Clean & Concentrator Kit (R1018; Zymo Research) to remove enzymes, free nucleotides, and other reaction components. The sgRNAs were eluted in nuclease-free water. RNA concentration and purity were assessed spectrophotometrically. The integrity of the synthesized sgRNAs were verified by denaturing 10% Tris–boric acid–EDTA–urea polyacrylamide gel electrophoresis (TBE-Urea PAGE; Invitrogen, Cat. No. EC68755BOX). Samples were mixed with RNA loading dye, heat-denatured at 70°C for 10 min, and resolved on the gel.

### Generation of Polq hemizygous (*Polq*^+/−^) clones

To generate *Polq*^+/−^ clones, one allele of the mouse *Polq* locus (~90 kb) was deleted using CRISPR/Cas9 ribonucleoprotein (RNP) approach. Two sgRNAs (5’-TCTCCCGCGCGGGAGGCCTA-3′; 5’-GTCCTTCAGCTCTCCCCAGC-3′) flanking the *Polq* locus were used for deletion of one allele of *Polq*. Cas9–sgRNA RNP complexes were preassembled by incubating EnGen Spy Cas9 NLS (New England Biolabs, Cat. No. M0646) with purified sgRNAs in Opti-MEM (Thermo Fisher Scientific, Cat. No. 31985062) to a final concentration of 3 µM. Founder conditional *Xrcc4* mES cells were transfected with the preassembled RNP complexes using Lipofectamine 2000 Transfection Reagent (Invitrogen, Cat. No. 11668019). 48 hours post-transfection, cells were trypsinized and plated at low density (250, 500, or 1000 cells per 6-cm dish) onto MEF feeder layers to allow clonal expansion. After 6 days, well-isolated single colonies were manually picked and transferred to MEF-coated 24-well plates (96 clones total). Following 4 days of expansion, each clone was split: half was cryopreserved in freezing medium and stored at −80°C, and the remaining half was seeded onto gelatin-coated 6-well plates for genomic DNA extraction. Deletion screening was performed by PCR using primers flanking the sgRNA target sites (5’-ACCGCACAAATTACGCTCTACC-3′; 5’-AGCAGCTCCACTCCTTCACATT-3′). PCR-positive clones were validated by Sanger sequencing to confirm deletion junctions. The remaining allele was independently amplified using primers targeting the 5′ (5’-AGGGCAACTCATTAGCTTCCGA-3′; 5’-AAGAGCCTGCATCCTACACCAC-3′) and 3′ (5’-GACCGGTTCTTTGCCATGTACG-3′; 5’-AGCAGCTCCACTCCTTCACATT-3′) flanking regions and sequenced to confirm absence of unintended mutations at the sgRNA target sites. Although two rounds of targeting were performed, no clones harboring a clean *Polq* deletion on one allele and an intact wild-type allele were obtained. However, clones carrying out-of-frame mutations were recovered. One clone (#9) containing a 10-bp out-of-frame deletion at the 3′ end of the *Polq* open reading frame (ORF) was revived from frozen stock and used for subsequent targeting strategies.

### Tagging of Polq with degron TAG

Tagging of Polq with degron TAG (dTAG-8×HA-SMASh-neomycin^R^ cassette) was performed using CRISPR/Cas9-mediated strategy in the Polq^+/−^ cell clone (#9) carrying the 10 bp deletion noted above. This approach enabled restoration of the wild type reading frame of the disrupted allele while simultaneously introducing a C-terminal degron-tagged fusion. Cas9–sgRNA ribonucleoprotein complexes targeting at the 3′ end of *Polq* ORF were preassembled in vitro. The RNPs were co-transfected into the mutant *Polq*^+/−^ mES cells together with 90 ng of a donor targeting construct using Lipofectamine 2000. The PvuI linearized donor targeting construct contained about 500 bp 5′ homology arm upstream of the Cas9 cut site, 500 bp 3′ homology arm downstream of the cut site and an in-frame C-terminal tagging cassette. The homology arms were complementary to genomic sequences flanking the Cas9-induced double-strand break at the 3′ end of the *Polq* ORF (sgRNA; 5’-CTCTTTTCTTTATTTGCAGG-3′), enabling precise HR-mediated integration. To prevent Cas9-mediated cutting of the donor template, silent mutations were incorporated within the protospacer adjacent motif (PAM) sequence. Four hours after transfection, four technical replicates (totaling 6.4 × 10^5^ cells) were pooled and plated onto neomycin-resistant MEF feeder-coated 6-cm dishes (four plates). Twenty-four hours post-transfection, culture medium was supplemented with 800 µg/mL G418 to select for neomycin-resistant clones. Selection was maintained for approximately one week, during which resistant colonies emerged. After one week of G418 selection, well-isolated single cell colonies were manually picked and transferred onto MEFs in four 24-well plates (96 clones total). Following 4 days of expansion, each clone was split into two fractions, half was cryopreserved in freezing medium and stored at −80°C. These stocks were preserved for recovery after genotyping confirmation. The remaining half was seeded onto gelatin-coated 6-well plates for genomic DNA. After 3 additional days of growth, genomic DNA was extracted from each clone. Correctly targeted clones were identified by PCR screening using primer sets designed to detect proper 5′ (5’-CCCATGCCTCTCAGAGTGAAAGA-3′; 5’-TTCTTCCCAGCCTCGGATCACC-3′) and 3′ (5’-CGCATCGCCTTCTATCGCCTTC-3′; 5’-TTCCTGGCCTGGCAGTGTTAGG-3′) junction integration. Clones positive for targeted integration were further validated by Sanger sequencing, enabling confirmation that targeted clones had restored a WT 3’ sequence in place of the previous 10 bp deletion. Clones with accurate cassette integration and WT *Polq* sequence were selected as hemizygous *Polq*^dTAG +/–^ clones for further experiments.

### Generation of Polq E688R point mutant clones

The *Polq*^E688R/–^ point mutant clones were generated by CRISPR/Cas9-mediated HR in the Polq^dTAG-8xHA/–^ cells. Briefly, a linear dsDNA donor template was designed to introduce a missense mutation at codon 688 of mouse *Polq*, converting glutamic acid (GAA) to arginine (AGA). The donor template contained ~500 bp 5’- and 3′-homology arms flanking the sgRNA-directed Cas9 cleavage site cloned using BamHI and EcoRI restriction sites in pUC19 (Addgene, plasmid #50005). To prevent cutting of the HR donor template, silent mutations were incorporated within the target PAM sequence. The sgRNA (5’-GGTGGCAGAGCTGGTTGGAG-3′) was selected such that Cas9 cleavage occurred approximately 8 bp from the intended mutation site. Cas9–sgRNA RNP complexes were preassembled in vitro at a final concentration of 3 µM. *Polq*^dTAG-8xHA/–^ cells were co-transfected with the preassembled RNP complex and 90 ng of linear donor plasmid using Lipofectamine 2000. 72 hours post-transfection, cells were trypsinized and plated at low density onto 6-cm dishes containing MEF feeder cells without antibiotic selection. Colonies were allowed to expand, and individual clones were manually picked between 6-7 days after plating. Genomic DNA was extracted from expanded clones and screened by PCR using primers flanking the edited region (5’-GCAGAGCTTGTTGGAGTTGAGAG-3′; 5’-CAAGACTCAGACCATGTGCACAC-3′; 5’-ATTAAATGCCTGCACCACCACG-3′; 5’-ACATCGAGCCAAGAACCCTCT-3′; 5’-CCACCTGCCCTTCTCTTCTTCC-3′; 5’-GGCACAAATAATCCAAGCACAGAG-3′). PCR amplicons were subjected to Sanger sequencing to confirm E688R (GAA→AGA) substitution. Correctly edited clones were designated *Polq*^dTag E688R/−^.

### Microhomology-mediated end joining (MMEJ) FACS assays

Asynchronously growing mES cells were trypsinized and reseeded 24 h prior to transfection to ensure optimal proliferation and cell density. Immediately before transfection, cells were detached using 0.25% trypsin and neutralized with serum-containing medium. Cells were counted using hemocytometer and maintained in suspension, and each clone was divided equally into two sterile microcentrifuge tubes corresponding to treatment conditions. One group received 1 µL DMSO per mL of culture medium as a vehicle control, while the second group was treated with 1 µL of 3 mM ART558 per mL. For each condition, 1.6 × 10^5^ cells suspended in 200 µL medium were seeded into individual wells of a 24-well plate and transfected using Lipofectamine 2000. The following plasmid combinations were used: (i) 0.5 µg pcDNA3β empty vector alone (background GFP^+^%; one cell line per experiment); (ii) 0.475 µg pcDNA3β empty vector plus 25 ng mCherry plasmid (transfection effeciency control; one cell line per experiment); and (iii) 0.45 µg pcDNA3β empty vector plus 25 ng mCherry plasmid and 25 ng linear GFP_MMEJ6 plasmid. The total amount of transfected DNA was kept constant across all conditions. Six hours after transfection, 1 mL of complete mES medium was added to each well along with either 1 µL DMSO or 1 µL of 3 mM ART558 to maintain the respective treatment conditions. Cells were returned to the incubator and cultured for an additional 18 h. At 24 h post-transfection, cells were harvested and analyzed by flow cytometry. The MMEJ frequency was calculated as the ratio of GFP^+^RFP^+^: Total RFP^+^, following background subtraction, expressed as a percentage (Fig. S11A). These MMEJ values were compared between DMSO- and ART558-treated groups within each clone and the level of TMEJ was calculated by subtracting the MMEJ frequency in the ART558-treated sample from the MMEJ frequency in the DMSO-treated sample (Fig. 5D).

### Western blotting

Freshly harvested cells were washed once with 1× PBS, pelleted by centrifugation, and lysed in RIPA lysis and extraction buffer (Thermo Fisher Scientific, Cat. No. 89901) mixed with 1× Halt Protease Inhibitor Cocktail (Thermo Fisher Scientific, Cat. No. 78438). Cell lysates were incubated on ice for 30 minutes with intermittent vortexing every 5 minutes to ensure efficient lysis. Samples were then sonicated using a probe sonicator (Heat Systems Misonix Sonicator Ultrasonic Processor XL) at 4°C for 10 seconds at 30% amplitude, repeated twice. Following sonication, lysates were centrifuged at 14,000 rpm for 30 minutes at 4°C to pellet cellular debris. The supernatant was transferred to fresh microcentrifuge tubes and stored at −80°C until further use. Total protein concentration was determined using the Pierce Rapid Gold BCA Protein Assay Kit (Thermo Fisher Scientific, Cat. No. A53226) according to the manufacturer’s instructions. Absorbance was measured at 480 nm using Spectramax Mini microplate reader. Based on the quantified protein concentration, 30 µg of total protein from each sample was prepared for electrophoresis. Samples were mixed with LDS sample buffer (Thermo Fisher Scientific, Cat. No. NP0007), sample reducing agent (Thermo Fisher Scientific, Cat. No. B0009) and RIPA lysis buffer to a final volume of 28 µL. The samples were denatured by boiling at 95°C for 10 minutes. Proteins were separated on 3– 8% Tris-Acetate SDS–PAGE gel (NuPAGE 3–8% Tris-Acetate Mini protein gels; Thermo Fisher Scientific, Cat. No. EA03785BOX) and electrophoresed at 50 V for 20 hours at 4°C. Following electrophoresis, proteins were transferred onto nitrocellulose membrane (Invitrogen, Cat. No. LC2001) at 100 mA for 6 hours at 4°C. Protein transfer and equal loading were verified by Ponceau staining and imaging. Membranes were blocked in 5% nonfat milk (in 1x PBST) for 1 hour at room temperature with gentle rocking. The membranes were then incubated overnight at 4°C with primary antibodies diluted in 5% skim milk: anti-HA (1:1000; Abcam, Cat No. ab9110) and anti-Vinculin (1:2500; Abcam, ab129002). After primary antibody incubation, membranes were washed in 1x PBST and incubated with HRP-conjugated anti-rabbit secondary antibody (1:5000 dilution in 5% skim milk) for 1 hour at room temperature with gentle rocking. Protein bands were visualized using chemiluminescent detection and imaged on Amersham and BioRad ChemiDoc Imagers.

### Polθ-H-RPA on MH ssDNA complex preparation for cryo-EM

Polθ-H and RPA were buffer exchanged into cryo-EM buffer (10 mM Na·HEPES pH 7.4 at 22 °C, 150 mM NaCl, 0.5 mM TCEP) by five rounds of successive dilution and concentration in Vivaspin 500 centrifugal filters (30-kDa MWCO, Sartorius) prior to assembly. MH ssDNA (TTTTTTTTTTTTTTTTCACTGTGAGCTTAGCGTTAGAGTAGGTTTTTTTGCCCGGG) was resuspended in cryo-EM buffer. DNA and RPA were combined with cryo-EM buffer and incubated at 4°C for 30 min. Polθ-H, AMP-PNP, and MgCl_2_ were added and the reaction was incubated at 4°C overnight. Final 30µL reactions contained 8 µM Polθ-H, 10.7 µM RPA, 5.3 µM DNA, 2 mM AMP-PNP and 5 mM MgCl_2_. Cryo-EM grids (UltrAuFoil R 2/2, 200 Mesh, Quantifoil Micro Tools GmbH) were glow discharged for 300 s at 30 mA with a Pelco Easiglow plasma discharge system. 3 µL of sample was applied to the grid. After incubation for 8s, each grid was blotted for 3s at blot force 8. The grid was subsequently vitrified by plunging into liquid ethane using a Vitrobot Mark IV (FEI) operating at 4 °C and 100 % humidity.

### Cryo-EM data collection

Cryo-EM data was collected on a Thermo Fisher Scientific Titan Krios operating at 300 keV equipped with a Falcon 4i direct electron detector and Selectris energy filter. Data collection was automated using Thermo Scientific Smart EPU software. 9,741 micrographs were collected with a pixel size of 0.94 Å across a defocus range of 1-1.8 µm. The dataset was collected with 48 movie frames at an exposure time of 5.62 s and electron flux of 9.07 e^-^/Å^2^ per second resulting in a total exposure dose of 50.94 e^-^/Å^2^

### Cryo-EM data analysis

Initial image processing was performed in CryoSPARC (v.4.7.1). Movies were aligned using CryoSPARC Live Patch Motion Correction followed by Patch CTF Estimation. CryoSPARC blob picker was used to pick 5,092,152 particles. Particles were extracted to a box size of 300^2^ pixels. All classification and refinement were done in CryoSPARC. Initial 2D classification was used to identify globular protein particles, which were subsequently used for *ab initio* reconstruction of a Polθ-H-RPA-ssDNA complex. Junk particles were removed by three rounds of heterogeneous refinement resulting in 1,090,623 particles. Non-uniform refinement of these particles resulted in a 3.4 Å map of the Polθ-H-RPA dimer. To reduce map anisotropy and resolve compositional heterogeneity, these particles were used as input for alignment-free 3-D classification, using 5 classes, an 8 Å filter resolution, and PCA initialization. 231,659 particles from the best resolved class were selected for further non-uniform refinement resulting in 3.4 Å map of reduced anisotropy (Map A). To improve local map quality, each individual subunit of the Polθ-H-RPA dimer was subjected to local refinement. Soft masks encompassing the region of interest were generated and local refinements were conducted independently using a rotation search distance of 5° and shift search extent of 5 Å. This resulted in 3.4 Å maps of the two RPA molecules (Maps D and E, respectively) and two 3.2 Å maps of Polθ-H monomers (Maps B and C). Locally refined maps were combined into a composite reconstruction (Map F) using the PHENIX (v2.2.1-6174) combine focused refinements tool. The resulting map was then processed with DeepEMhancer resulting in a sharpened composite reconstruction (Map G).

### Model building and refinement

Two copies of RPA (PDB: 8RK2) and one Polθ-H dimer in a microhomology annealed state (PDB: 9BHA) were rigid-body docked into map F in UCSF ChimeraX. Stem-loop DNA in the starting model was removed due to large steric clashes with RPA. The model was real space refined in PHENIX with five macrocycles of global minimization, local grid search, and NHQ flips. Iterative rounds of manual adjustment in ISOLDE (v1.12.1) and Coot (v1) with real space refinement resulted in a final model with good geometry and density match.

### Structural model preparation

The amino acid sequences of RPA1, RPA2, and RPA3 were retrieved from UniProt (*57*), (RPA1: P27694; RPA2: P15927; RPA3: P35244). The RPA1– RPA2–RPA3–ssDNA (RPA123–ssDNA) complex was constructed using the available X-ray crystal structures (PDB IDs: 4GOP (*28*) and 8RK2 (*27*)) together with our RPA123-ssDNA-Polθ complex modeled using AlphaFold3 which had ssDNA threaded into Polθ helicase (Alphafold3 model). The ssDNA was modelled in contact with RPA1 and RPA2, using PDB structures 4GOP and 8RK2 (ssDNA not resolved but present) as structural references. Because RPA1 contains a zinc-binding domain, the associated Zn^2+^ ion was retained in the structural model. Protonation states of ionizable residues were assigned based on predictions from the H++ server (*58*). The helicase domain of DNA polymerase theta (Polθ; UniProt ID: O75417) was extracted from our AlphaFold3 model and protonated independently using the H++ server.

### Molecular dynamics simulations

The RPA123–ssDNA complex and the Polθ helicase domain were initially simulated independently using classical, unbiased, all-atom molecular dynamics (MD). Each system was solvated in a rectangular box of explicit TIP3P (*59*) water molecules extending at least 10 Å from the solute in all directions. Na^+^ and Cl^−^ ions were added to neutralize the net charge of each system and subsequently achieve a final salt concentration of 150 mM.

Protein residues were parameterized using the AMBER ff19SB ^(*60*)^ force field, whereas DNA was described using the OL21 (*61*) force field. The Zn^2+^ ion in RPA1, coordinated by Cys481, Cys486, Cys500, and Cys503, was modeled using the ZAFF (*62*) force field. Periodic boundary conditions were applied, and long-range electrostatic interactions were treated using the particle mesh Ewald (PME) (*63*) method. A 10 Å cutoff was applied to nonbonded interactions. Bonds involving hydrogen atoms were constrained using the SHAKE (*64*) algorithm, permitting an integration time step of 2 fs.

Each system was energy-minimized for 5,000 steps, consisting of 2,000 steps of steepest-descent minimization followed by 3,000 steps of conjugate-gradient minimization. The systems were then heated to 300 K over 100 ps under constant-volume and constant-temperature (NVT) conditions using a Langevin thermostat (*65*). Following heating, the systems were equilibrated for 200 ps under NVT conditions and subsequently for 10 ns under constant-pressure and constant-temperature (NPT) conditions. Production simulations were performed for 100 ns under NPT conditions at 300 K, with trajectory coordinates saved every 20 ps. All simulations were performed using AMBER24 (*66-68*) with the GPU-accelerated pmemd.cuda engine on NVIDIA GPUs.

### Construction and refinement of the Polθ–RPA123–ssDNA complex based on MD simulations

To construct the monomeric Polθ–RPA123–ssDNA complex, 20 conformations were sampled at 5 ns intervals from each of the independently simulated RPA123–ssDNA and Polθ helicase trajectories. Each sampled structure was superimposed onto the corresponding component of AlphaFold3 model using Cα atoms. Candidate Polθ– RPA123–ssDNA assemblies were generated from the aligned structures and evaluated for steric clashes, defined as overlaps between atomic van der Waals radii. Among the evaluated conformational combinations, the pair exhibiting the fewest inter-component clashes was selected to construct the initial monomeric complex.

Residual steric clashes introduced during assembly were relieved by staged energy minimization in explicit solvent. Positional restraints were applied to non-hydrogen heavy atoms of the protein and sequentially reduced using force constants of 100, 50, 20, and 10 kcal mol^−1^ Å^−2^. At each restraint level, 2,000 steps of steepest-descent minimization were followed by 3,000 steps of conjugate-gradient minimization. The system was subsequently subjected to an additional 5,000 steps of unrestrained energy minimization.

The refined monomeric complex was subsequently used to construct a Polθ–RPA123–ssDNA model with Polθ modelled as dimer. In addition to simulated structures, this modelling effort was guided by experimental X-ray crystal structures (PDB IDs: 4GOP (*28*), 8RK2 (*27*) and 8W0A (*69*)) and AlphaFold3 model as discussed previously. The monomeric and Polθ dimerized complex with RPA123–ssDNA systems were simulated using the same force fields, solvation conditions, electrostatic treatment, nonbonded cutoff, temperature-control method, and integration time step described above. For these assembled systems, however, a more gradual relaxation protocol was employed. Following staged minimization, each system was heated to 300 K over 100 ps under NVT conditions. This was followed by four successive restrained NVT equilibration stages of 50 ps each, during which positional restraints on protein non-hydrogen heavy atoms were progressively reduced using force constants of 100, 50, 20, and 10 kcal mol^−1^ Å^−2^. The systems were then equilibrated without positional restraints for 20 ns. The final structures obtained from these simulations were used as the equilibrated monomeric and dimeric Polθ–RPA123–ssDNA complex models (Fig. S9).

## Supplementary figures

**Figure S1.**
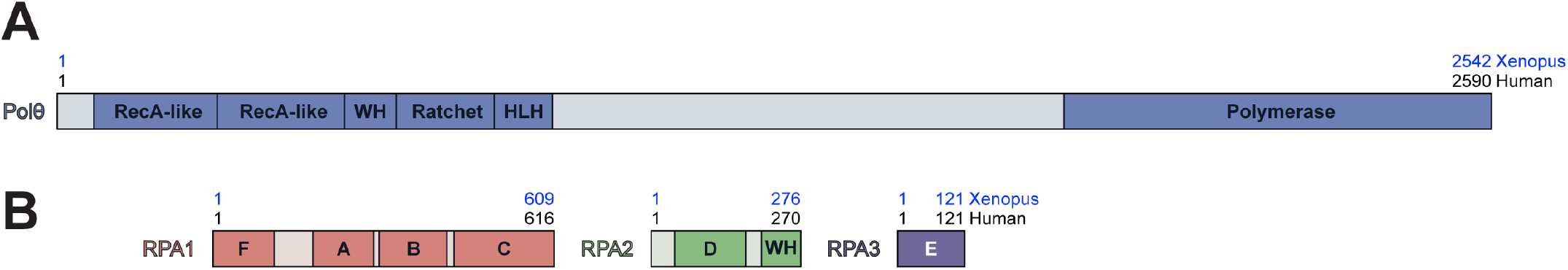
Domain organization of Polθ and RPA. **(A)** Scheme of Polθ. Black and blue numbers refer to human and *Xenopus laevis* residues, respectively. **(B)** Same as (A) for RPA1, RPA2, and RPA3 with DNA-binding domains (DBDs) A-F noted. WH = Winged Helix. HLH = helix-loop-helix.

**Figure S2.**
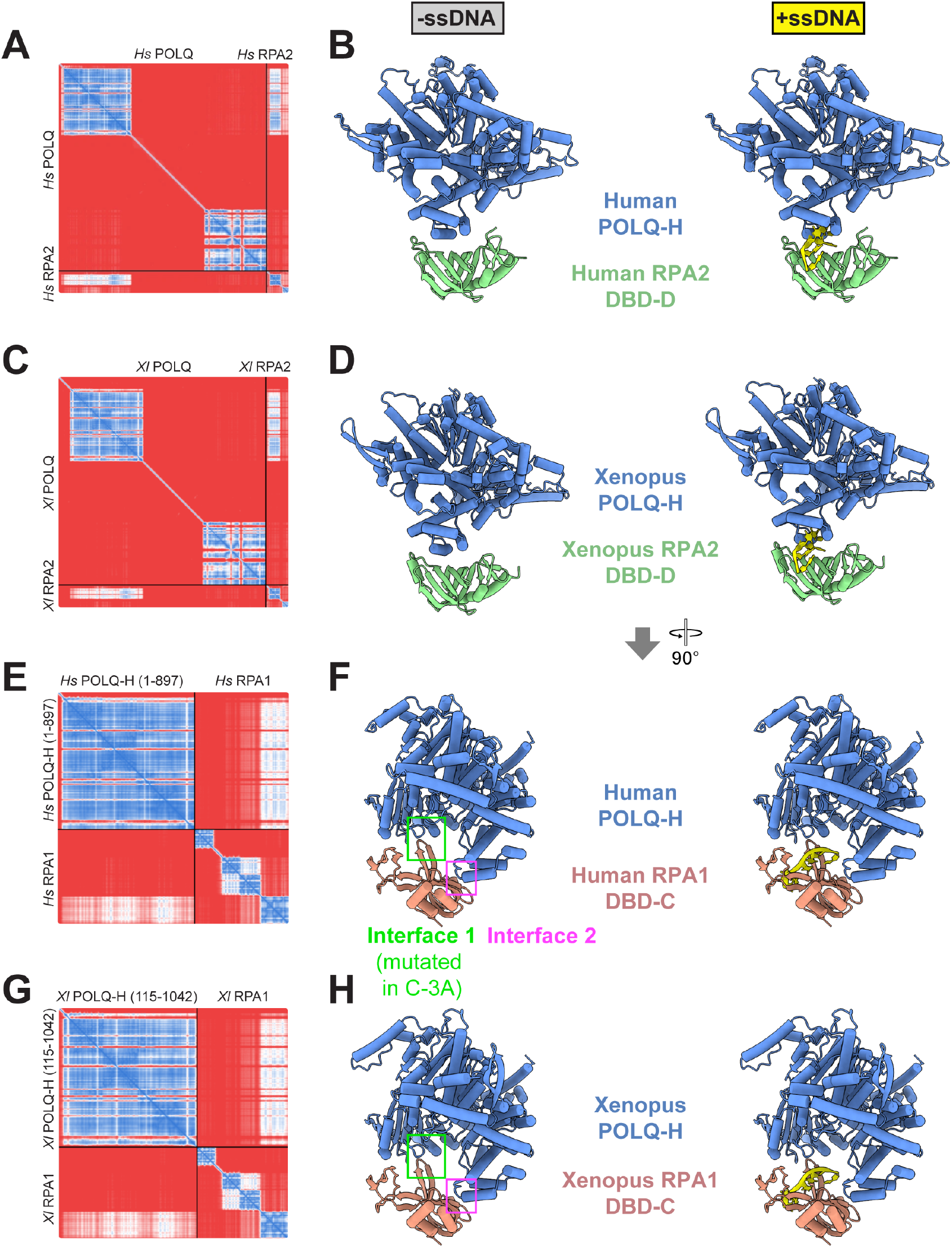
AlphaFold predictions between Polθ and RPA1 and RPA2. **(A)** PAE plot of the best model for the interaction between human Polθ and human RPA2 (downloaded from predictomes.org). **(B)** Predicted structure of human Polθ (only residues 37-897 comprising the helicase domain are shown) and human RPA2 (only residues 43-171 comprising DBD-D are shown) without (left) and with (right) ssDNA, which was modeled by aligning the X-ray crystal structure of a fungal RPA-ssDNA complex (PDB:4GOP) with the DBD-D domain of human RPA2. Note that ssDNA clashes with the DBD-D-Polθ interaction. **(C)** PAE plot of the best model for the interaction between *Xenopus* Polθ and *Xenopus* RPA2. **(D)** Predicted structure of *Xenopus* Polθ (only residues 150-1027 comprising the helicase domain are shown) and *Xenopus* RPA2 (only residues 48-175 comprising DBD-D are shown) without (left) and with (right) ssDNA aligned to DBD-D as in (B). ssDNA clashes with the DBD-D-Polθ interaction. **(E)** PAE plot for best model of interaction between human Polθ-H (residues 1-897) and human RPA1. **(F)** Predicted structure of Human Polθ-H (residues 37-897 shown) with human RPA1 (only residues 32-171 are shown) without (left) and with (right) ssDNA modeled by aligning the X-ray crystal structure of fungal RPA-ssDNA complex (PDB:4GOP) with the DBD-C domain of human RPA1. ssDNA does not clash with the DBD-C-Polθ interaction. **(G)** PAE plot for the best model of interaction between *Xenopus* Polθ-H (residues 115-1042) and *Xenopus* RPA1. **(H)** Predicted structure of *Xenopus* Polθ-H (residues 150-1027 shown) with *Xenopus* RPA1 (residues 48-175 shown) without (left) and with (right) ssDNA modeled by aligning the RPA-ssDNA complex (PDB:4GOP) with the DBD-C domain of *Xenopus* RPA1.

**Figure S3.**
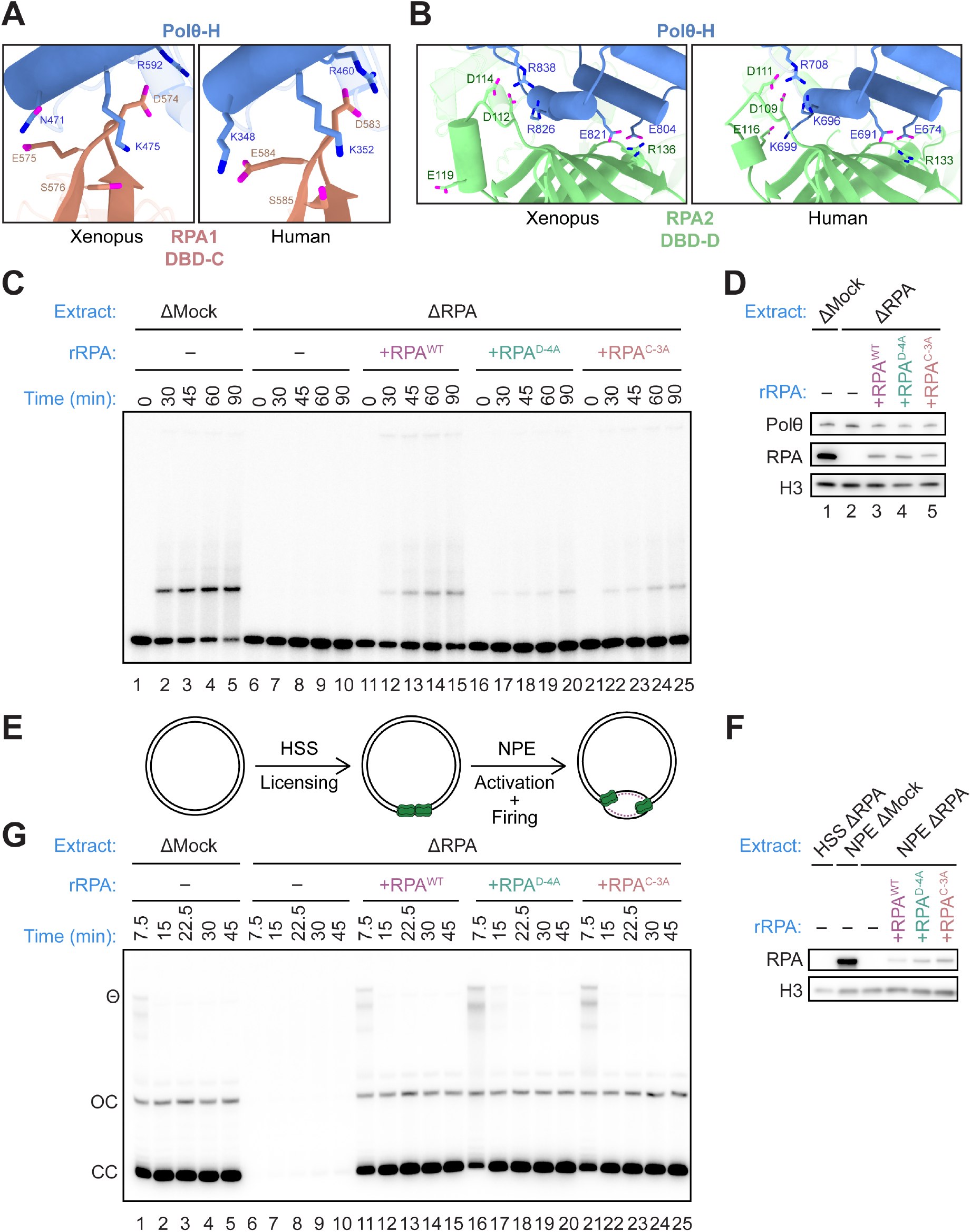
RPA mutant design and characterization. **(A)** Close up view of the predicted interface between Polθ-H and the RPA1 DBD-C domain in Xenopus (left) and human (right). The three residues that were mutated in *Xenopus* and human RPA, respectively, to generate RPA^C-3A^, as well as residues in Polθ-H they are predicted to interact with, are shown in stick form. **(B)** Same as (A), but showing RPA residues mutated at the interface between Polθ-H and RPA2 DBD-D to generate *Xenopus* and human RPA^D-4A^, plus interacting residues in Polθ-H. **(C)** Representative gel of the TMEJ assay depicted in Figure 1F. **(D)** Western blot of extract used for a representative example of the TMEJ assay in Figure 1F and Fig. S3C. **(E)** *Xenopus* replication assay. A plasmid was incubated in RPA-depleted high-speed supernatant of total egg lysate in the presence of [α-^32^P]dATP to promote licensing, followed by addition of NPE that was mock-depleted or RPA-depleted and supplemented with the indicated RPA constructs to promote replication initiation. **(F)** Protein samples from the replication reaction were blotted for RPA and histone H3. **(G)** DNA replication was carried out as described in (E) with the indicated conditions. Results were visualized using native agarose gel electrophoresis followed by autoradiography. Representative autoradiograph is shown.

**Figure S4.**
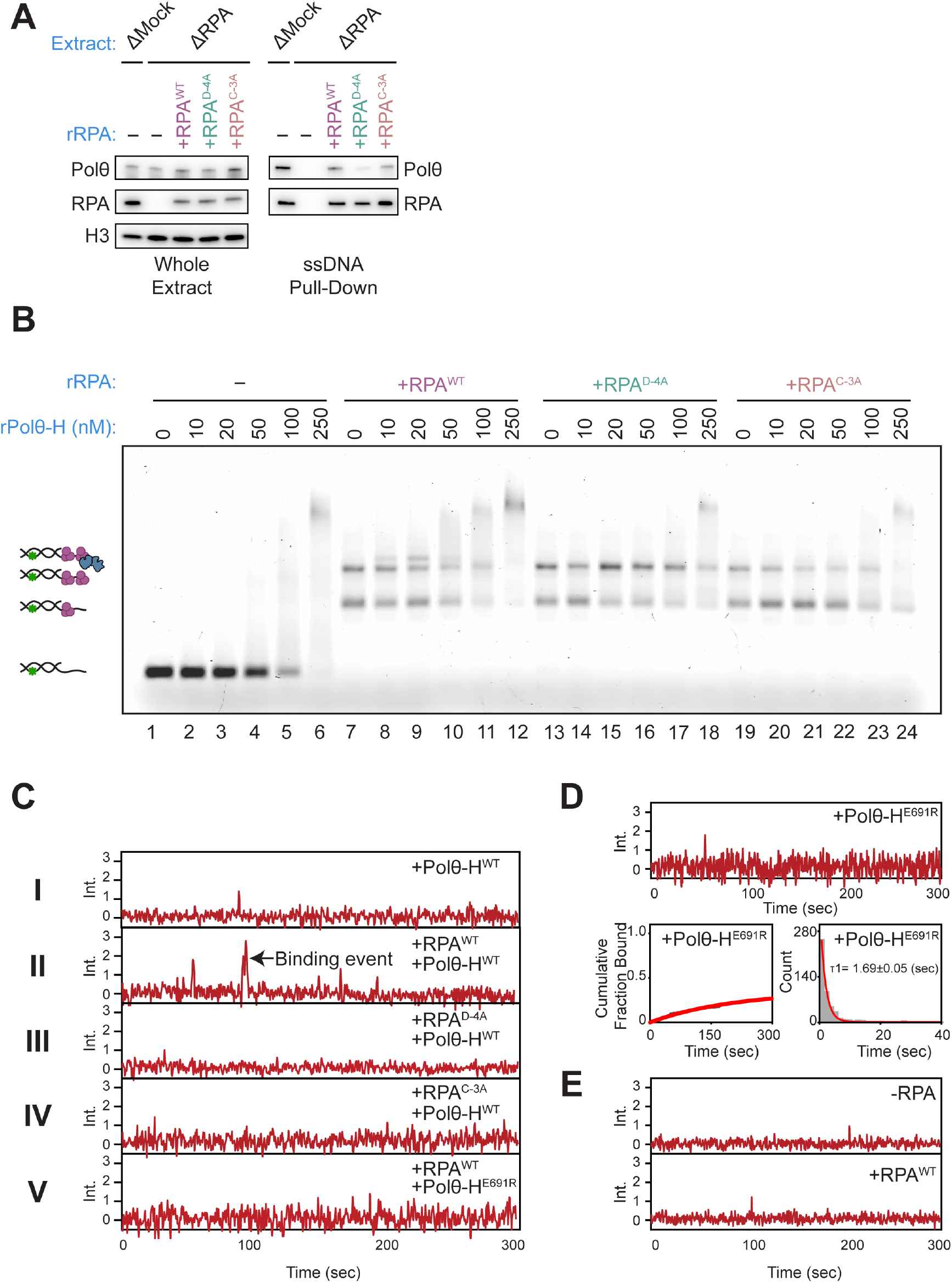
Polθ recruitment requires interactions with RPA. **(A)** Representative western blot of protein samples from the recruitment assay quantified in Figure 2B. **(B)** Representative gel shift quantified in Figure 2C. DNA containing a 3′ ssDNA overhang was incubated with increasing concentrations of hPolθ-H in the absence or presence of the indicated hRPA variants. DNA binding (%) was calculated as the signal corresponding to hPolθ-H-bound DNA divided by the total DNA signal in each lane, multiplied by 100. hRPA–DNA complexes lacking hPolθ-H were excluded from the bound fraction. **(C)** Representative Cy5 fluorescence intensity trajectories from the single-molecule colocalization assay illustrated in Figure 2D and analyzed in Figure 2E. Cy5-labeled hPolθ-H binding was monitored at the positions of immobilized Cy3-labeled 45-nt ssDNA substrates with a free 3′ end. Trajectories I–V correspond to the conditions shown in Figure 2E. The arrow marks an example binding event. **(D)** Binding of Cy5-labeled Polθ-H^E691R^ to immobilized Cy3-labeled DNA with a free 3’ end without RPA. *Top:* example trajectory, *Bottom left:* cumulative fraction bound and *Bottom right:* dwell time histogram. **(E)** Representative trajectories from the single-molecule colocalization assay shown in Figure 2G, using a 45 nt ssDNA substrate tethered at its 3′ end, in the absence (top) or presence (bottom) of RPA.

**Figure S5.**
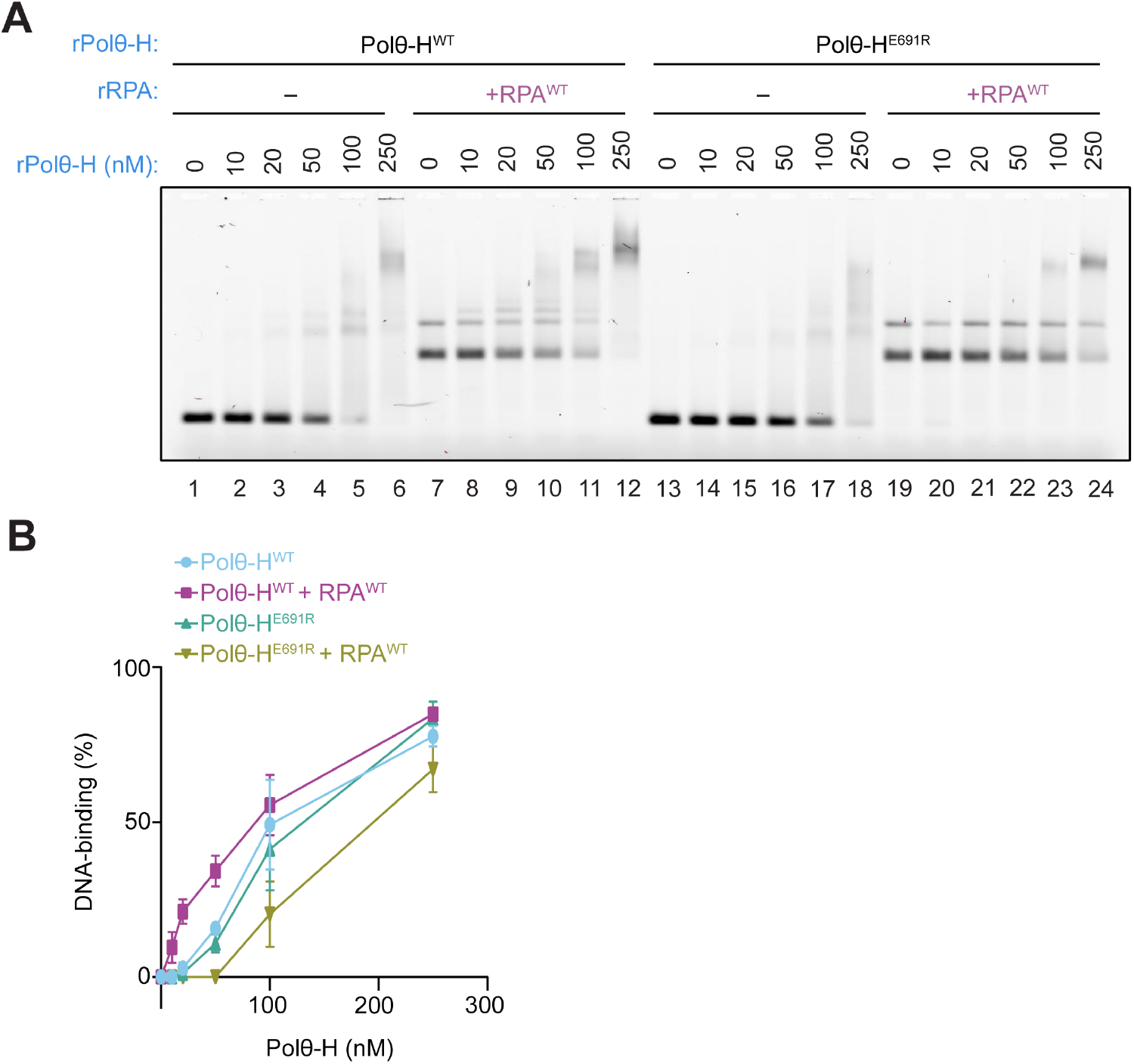
DNA binding by Polθ-H WT and E691R in the absence or presence of RPA. **(A)** Representative gel shift performed as in Figure S4B using the indicated conditions with hPolθ-H^WT^, hPolθ-H^E691R^, and hRPA^WT^. **(B)** Quantification of DNA binding in (A). DNA binding (%) was calculated as the signal corresponding to Polθ-H-bound DNA divided by the total DNA signal in each lane, multiplied by 100. RPA–DNA complexes lacking Polθ-H were excluded from the bound fraction. Data represent the mean ± SEM from three independent experiments.

**Figure S6.**
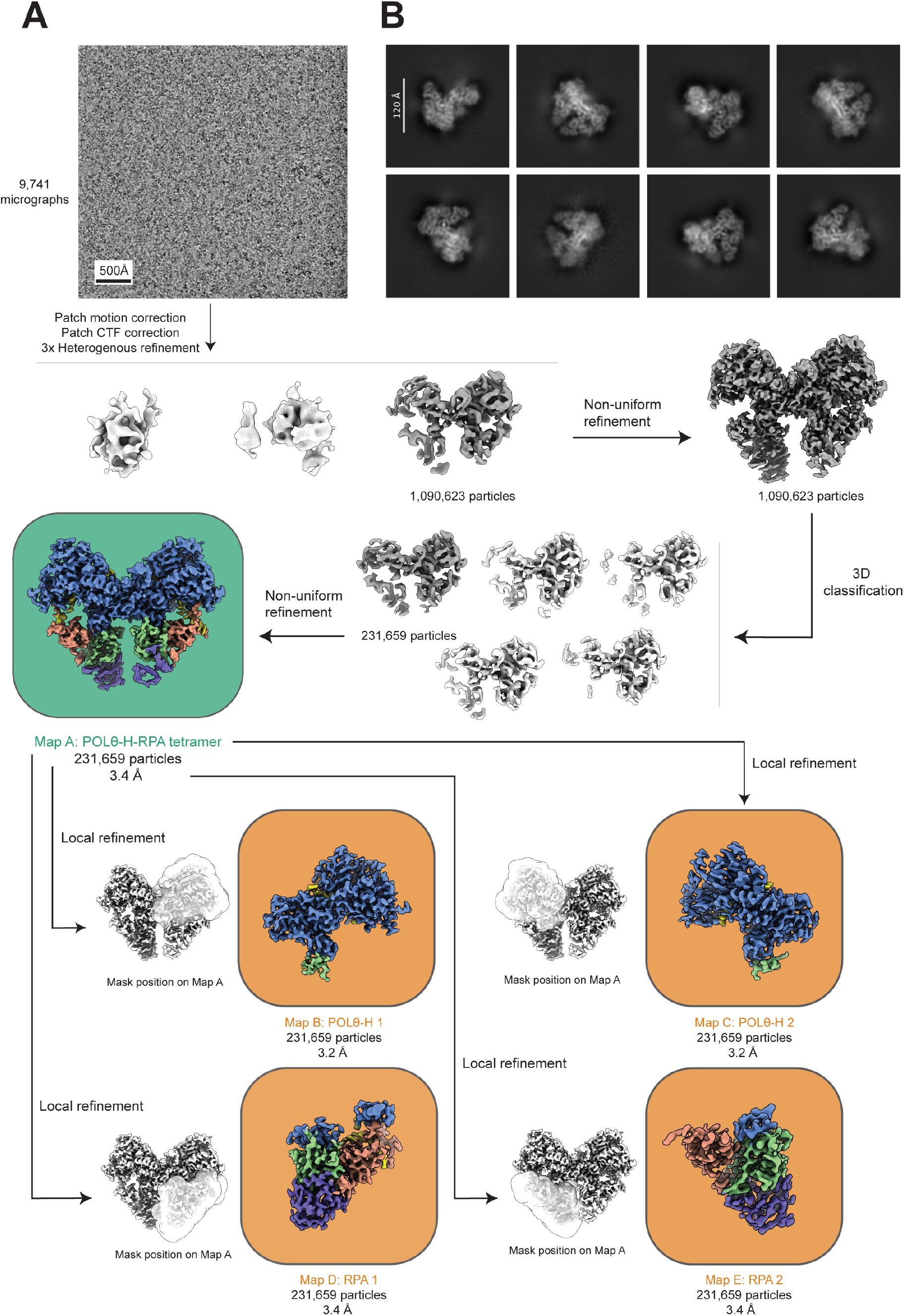
Cryo-EM data processing. **(A)** Workflow of image processing in single-particle cryo-EM analysis. See Methods for details. Scale bar: 500 Å. **(B)** Representative 2D classes of the Polθ-H-RPA dimer. Scale bar: 120 Å.

**Figure S7.**
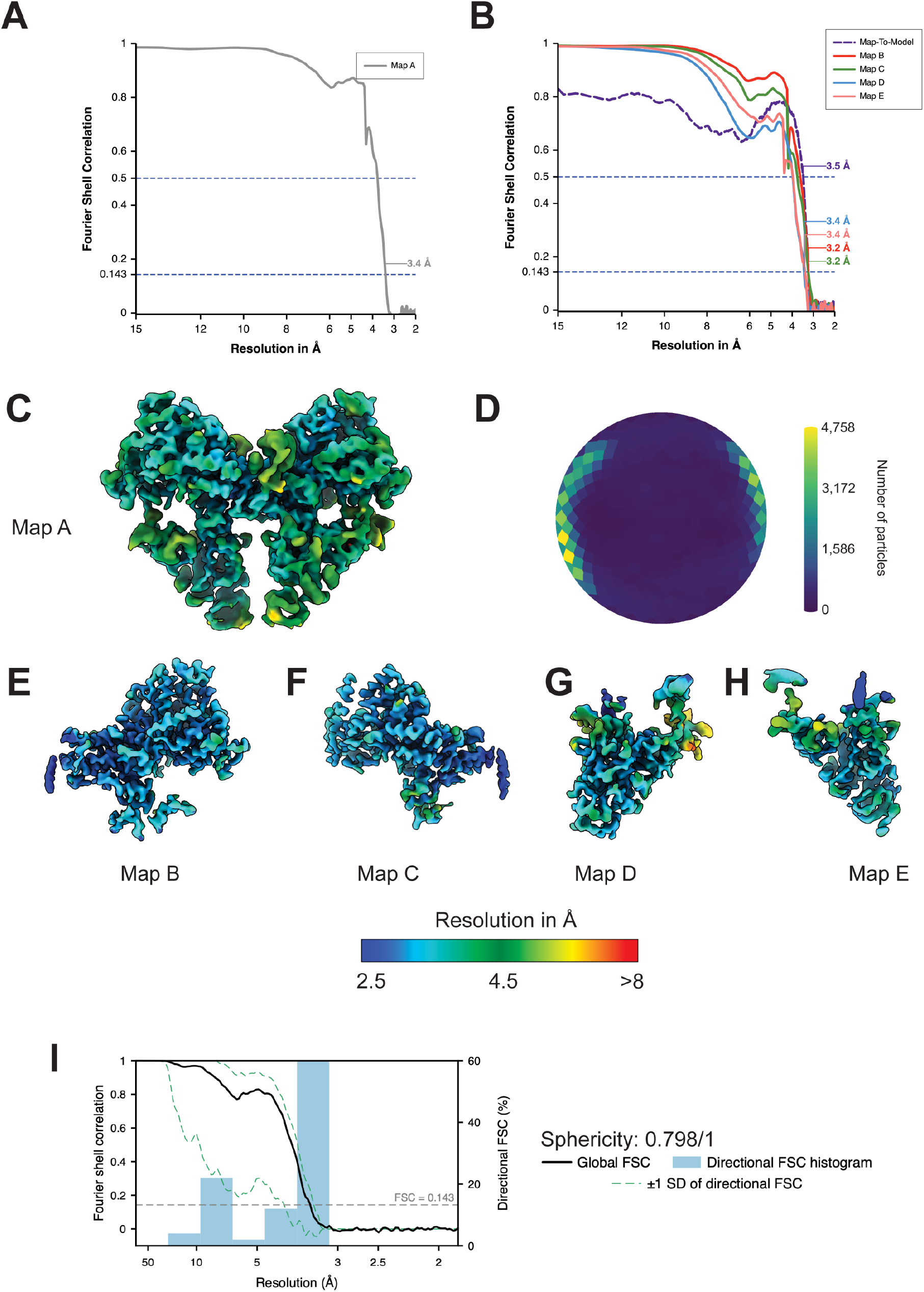
Cryo-EM data validation. **(A)** Fourier shell correlation (FSC) curve for the global Polθ-H-RPA dimer (map A). **(B)**. Fourier shell correlation curves for local refinement maps (maps B-E) and map-to-model FSC. **(C)** Local resolution map of the global Polθ-H-RPA dimer (map A). All maps are colored by local resolution as indicated. **(D)** Angular distribution plot of the global Polθ-H-RPA dimer (map A). **(E)** Local resolution map of Polθ-H copy 1 (map B). **(F)** Local resolution map of Polθ-H copy 2 (map C). **(G)** Local resolution map of RPA copy 1 (map D). **(H)** Local resolution map of RPA copy 2 (map E). **(I)** 3D Fourier shell correlation plot, directional Fourier shell correlation histogram and sphericity of the global Polθ-H-RPA dimer (map A).

**Figure S8.**
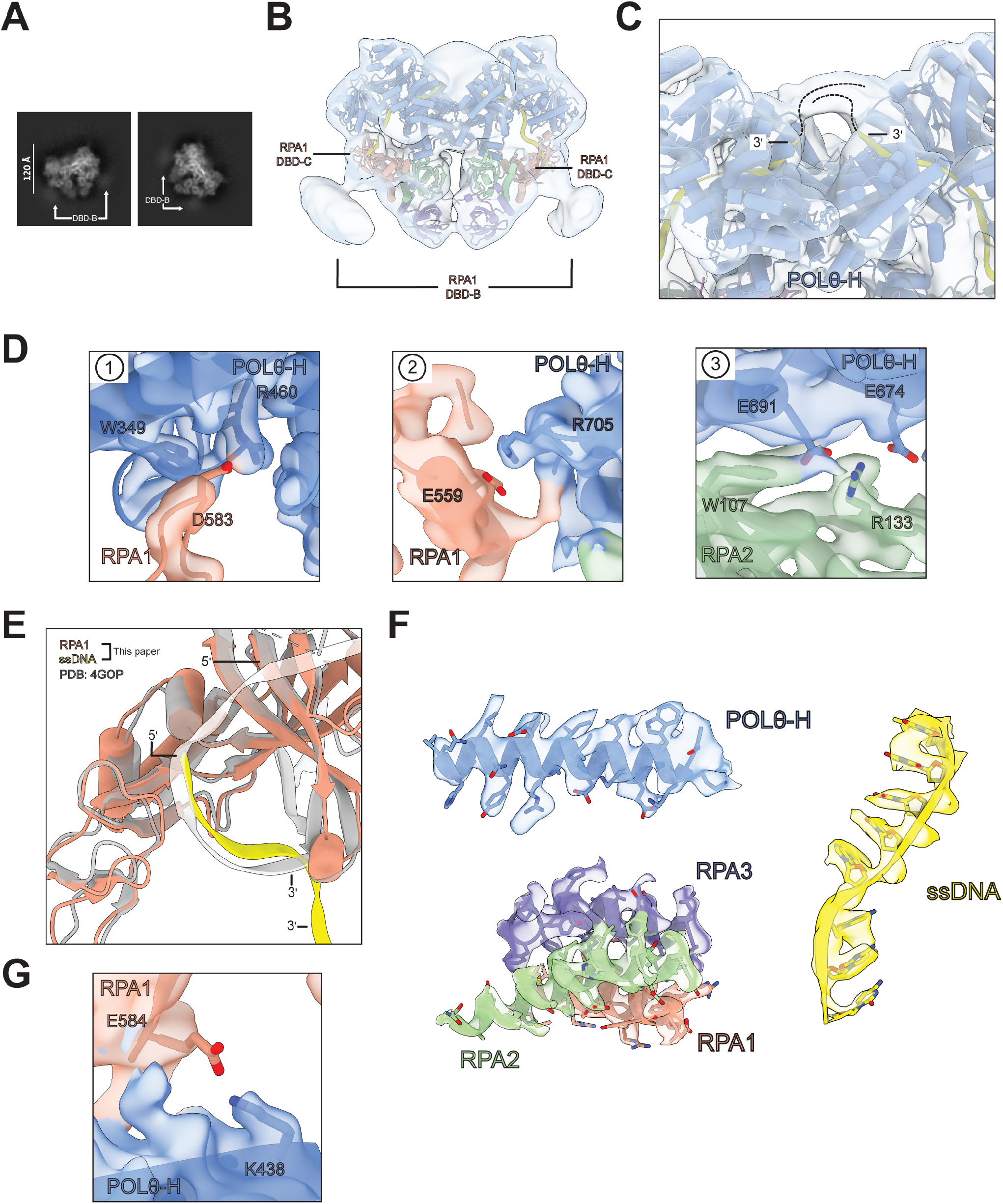
Structural details of the Polθ-H-RPA-ssDNA complex. **(A)** Representative 2D classes showing diffuse density consistent with RPA1 DBD-B. Scale bar: 120 Å. **(B)** Low-pass filtered map A with the atomic model docked reveals a low-resolution density adjacent to RPA1 DBD-C, consistent with flexibly tethered RPA1 DBD-B. **(C)** Low-pass-filtered map A with atomic model docked reveals low-resolution density consistent with microhomology annealed DNA in the Polθ-H dimer valley. **(D)** Reciprocal interfaces from the other protomer of the dimer, corresponding to those shown in Fig. 3C. **(E)** Overlay of the atomic models of RPA1 and ssDNA from this study and the *M. maydis* RPA-ssDNA complex (PDB: 4GOP). **(F)** Representative densities and atomic models of Polθ-H (Map B, residues 346-368), RPA1-3 (Map D, RPA1 residues 598-616, RPA2 residues 151-174, and RPA3 residues 95-114) and ssDNA (Map B, residues 46-53) **(G)** Interface of RPA1 E584 and Polθ-H K438 visible at high map contour.

**Figure S9.**
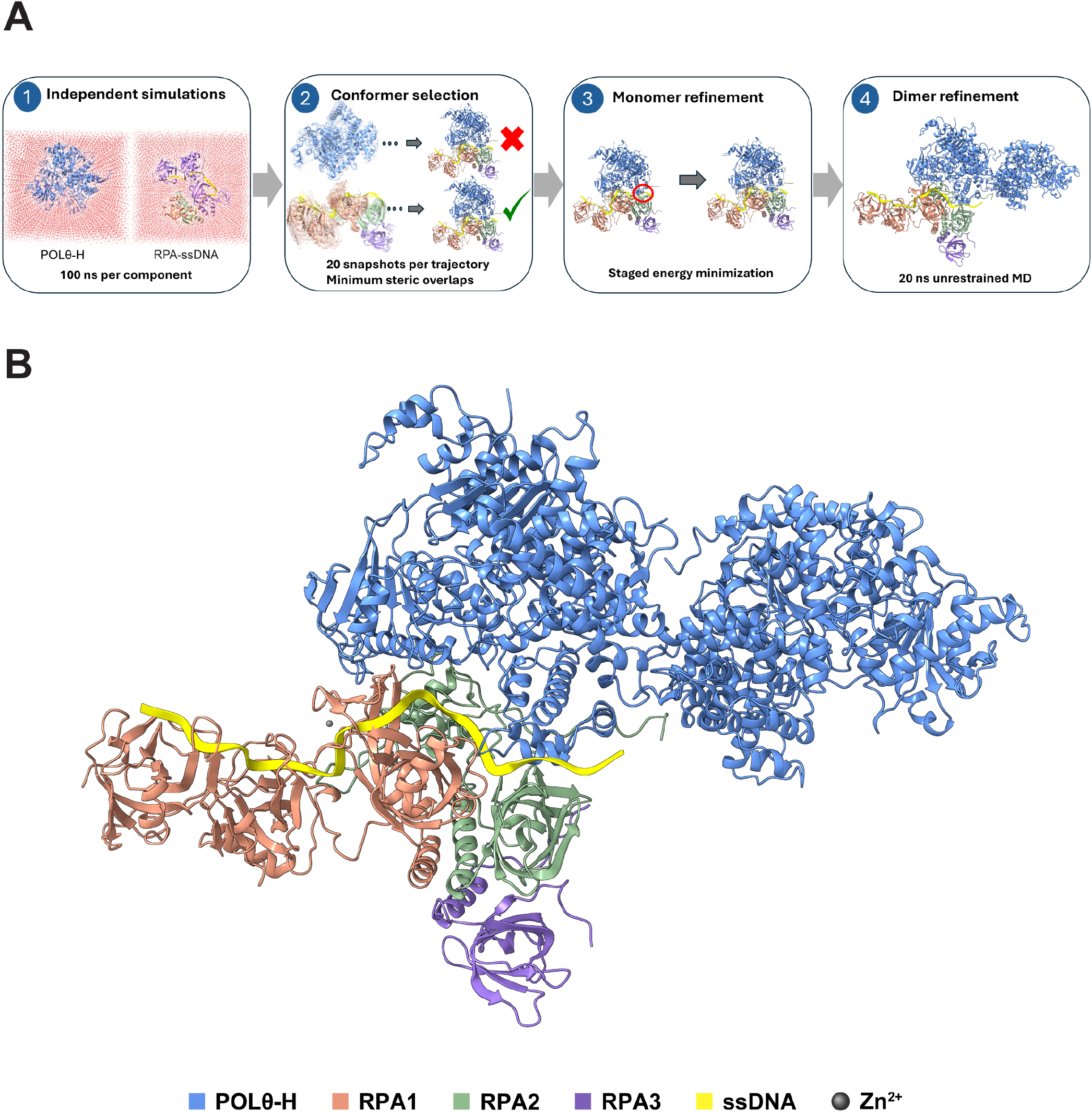
Molecular dynamics simulation of the Polθ helicase–RPA–ssDNA complex. **(A)** Molecular dynamics-guided assembly workflow. A multistep computational strategy was employed to identify sterically compatible configurations of Polθ helicase (Polθ-H) and ssDNA-bound replication protein A (RPA) and to construct a structural model of the resulting complex. Step 1, Independent simulations. Polθ-H and the RPA–ssDNA complex were subjected to independent 100 ns molecular dynamics (MD) simulations in explicit solvent to sample their conformational ensembles. The simulation boxes and surrounding water molecules are shown. Step 2, Conformer selection. Twenty snapshots were extracted from each trajectory and evaluated for their structural compatibility. Candidate assemblies were screened for steric overlaps between Polθ-H and RPA–ssDNA, with conformations exhibiting substantial clashes rejected (red cross) and sterically compatible configurations retained for further refinement (green check). Step 3, Monomer refinement. Selected configurations were subjected to staged energy minimization to alleviate local steric clashes and optimize the relative orientations of Polθ-H and RPA–ssDNA. The red circle highlights a region of steric overlap in the initial assembly that is resolved during refinement. Step 4, Dimer refinement. The refined assembly was incorporated into a dimeric Polθ-H model, followed by 20 ns of unrestrained MD simulation to allow conformational relaxation of the assembled complex. **(B)** Equilibrated structural model of the Polθ-H dimer bound to RPA–ssDNA. Representative structural model obtained following the MD-guided assembly and refinement procedure. The model illustrates a sterically compatible arrangement in which the Polθ-H dimer associates with RPA while RPA remains bound to ssDNA via DBD-C and DBD-D. Polθ-H is shown in blue, RPA1 in salmon, RPA2 in green, RPA3 in purple, ssDNA in yellow, and Zn^2+^ ion as gray spheres. The resulting configuration supports the structural feasibility of an intermediate in which Polθ-H can engage RPA while ssDNA remains associated with DBD-C and DBD-D.

**Figure S10.**
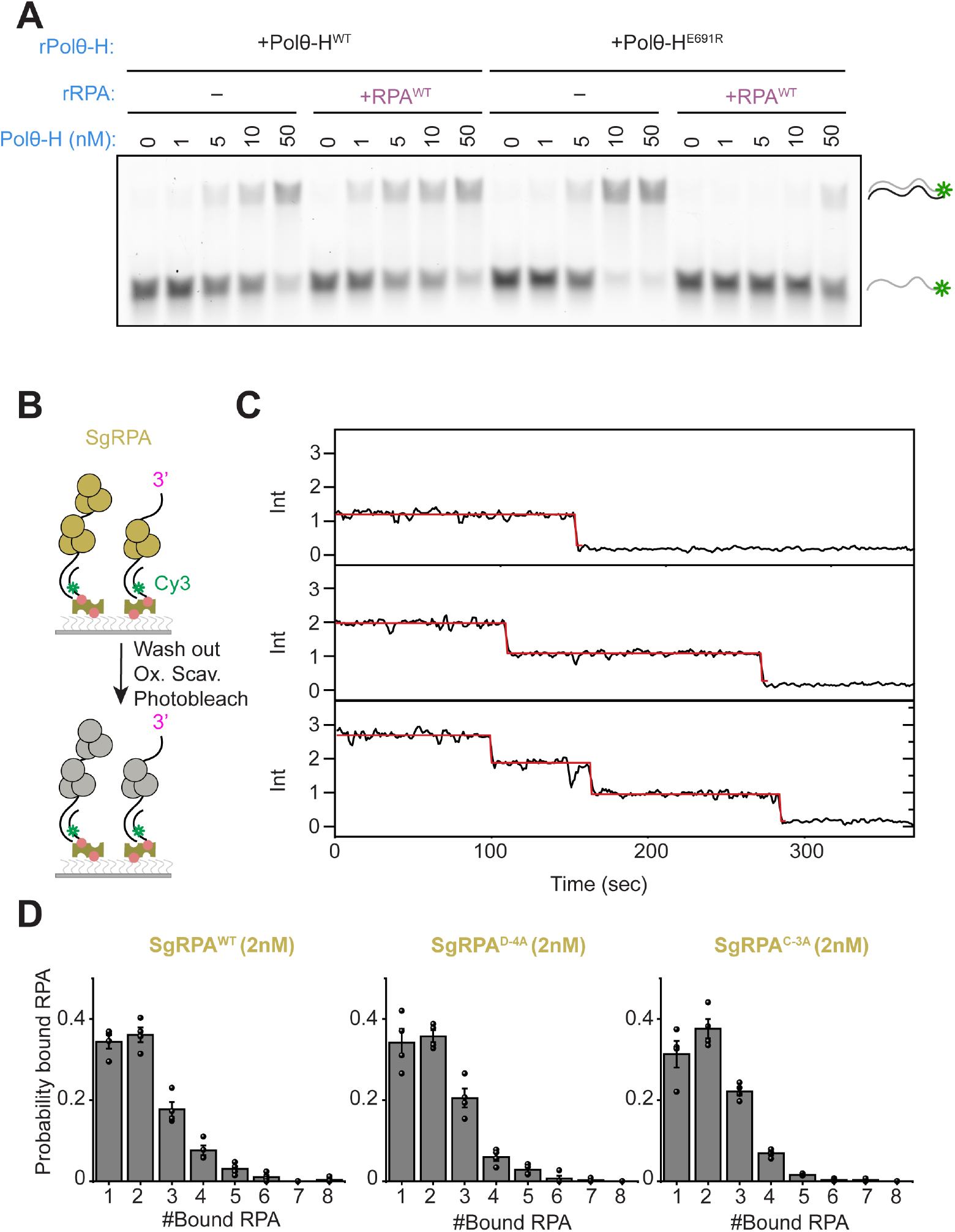
ssDNA annealing by Polθ-H and photobleaching analysis of DNA-bound SgRPA. **(A)** Representative ssDNA annealing assay quantified in Fig. 4D. **(B)** Schematic of the photobleaching assay using the same immobilized Cy3-labeled DNA substrate as in Figure 5A. The number of bound SgRPA molecules was determined from the number of photobleaching steps. **(C)** Example traces for SgRPA^WT^. Black lines show smoothed fluorescence trajectories, and red lines show stepwise idealized traces. From top to bottom: examples of 1, 2, and 3 photobleaching steps. **(D)** Histogram of the number of the indicated RPA molecules bound per DNA. Bars represent the mean ± SEM from four independent experiments (N=4). Each dot represents one independent experiment.

**Figure S11.**
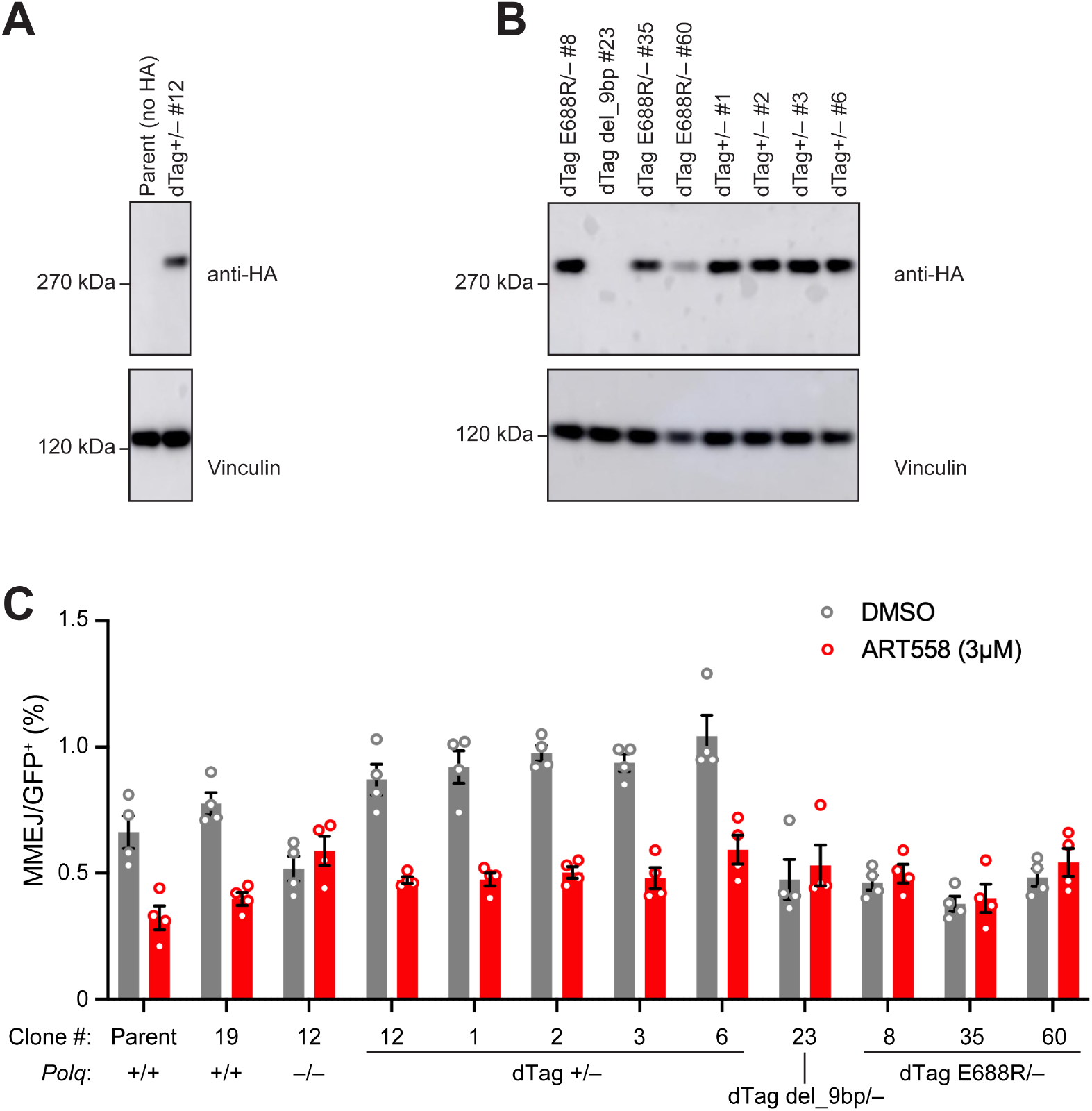
The Polθ-RPA interface is essential for TMEJ in cells. **(A)** Immunoblotting for anti-HA (upper panel) and vinculin (lower panel; loading control). Note anti-HA signal in *Polq*^dTag +/–^ cells and absence of anti-HA signal in parent *Xrcc4*^fl/fl^ cells, where Polθ is untagged. **(B)** Western blot showing abundance of HA-tagged Polθ (anti-HA, upper panel) in derivatives of *Polq*^dTag +/–^ clone #12. Vinculin: loading control (lower panel). HA-tagged Polθ abundance appears reduced in *Polq*^dTag E688R/–^ clone #60, in part due because the lane is underloaded. Note absence of HA signal in *Polq*^dTag del_9bp/–^ clone, likely reflecting instability of internally deleted gene product. **(C)** MMEJ products in assay shown in Figure 5D. MMEJ values shown here are GFP^+^RFP^+^ frequencies, normalized for transfection efficiency (Total RFP^+^) and expressed as a percentage. All *Polq* variants were derived from “Parent” *Xrcc4*^fl/fl^ clone shown. The last eight clones shown were all derived from founder *Polq*^dTag +/–^ clone #12. Mouse *Polq*^E688R^ corresponds to human *POLQ*^E691R^. Treatment with ART558 reduces MMEJ to a similar extent in parental *Polq*^+/+^ and *Polq*^dTag +/–^ cells. In contrast, in *Polq*^dTag E688R/–^ cells, basal MMEJ (DMSO condition) is reduced in comparison to cells expressing WT *Polq*, and ART558 addition has no effect on this basal level, showing that TMEJ is abolished in *Polq*^dTag E688R/–^ cells. Unpaired Welch’s *t*-test of DMSO *vs*. ART558 treatment in each clone: Parent P=0.0067; *Polq*^+/+^ p=0.0008; *Polq*^−/−^ p=0.395—not significant (NS); *Polq*^dTag +/–^ #12: p=0.0061; *Polq*^dTag +/–^ #1: p=0.033; *Polq*^dTag +/–^ #2: p<0.0001; *Polq*^dTag +/–^ #3: p=0.0002; *Polq*^dTag +/–^ #6: p=0.0056; *Polq*^dTag del_9bp/–^ #23: p=0.64 NS; *Polq*^dTag E688R/–^ #8: p=0.49 NS; *Polq*^dTag E688R/–^ #35: p=0.74 NS; *Polq*^dTag E688R/–^ #60: p=0.40 NS.

**Table S1.** Single-exponential analysis of Polθ-H dwell-time distributions under the indicated RPA and DNA-substrate conditions.

| Panel | Condition | Number of Dwell events | Decay constant, $\tau$ (s) | Fit SE (s) |
| --- | --- | --- | --- | --- |
| Fig. 2E | -RPA/ Polθ-H<sup>WT</sup> | 406 | 1.24072 | 0.01511 |
| Fig. 2E | +RPA<sup>WT</sup>/ Polθ-H<sup>WT</sup> | 3177 | 1.47776 | 0.02897 |
| Fig. 2E | +RPA<sup>D-4A</sup>/ Polθ-H<sup>WT</sup> | 100 | 0.93394 | 0.02104 |
| Fig. 2E | +RPA<sup>C-3A</sup>/ Polθ-H<sup>WT</sup> | 271 | 0.91889 | 0.00611 |
| Fig. 2E | +RPA<sup>WT</sup>/ Polθ-H<sup>E691R</sup> | 426 | 1.60984 | 0.06109 |
| Fig. 2G | -RPA/ Polθ-H<sup>WT</sup> (5'end) | 327 | 0.86519 | 0.02312 |
| Fig. 2G | +RPA<sup>WT</sup>/ Polθ-H<sup>WT</sup> (5'end) | 180 | 0.97809 | 0.02447 |
| Fig. S4D | -RPA/ Polθ-H<sup>E691R</sup> | 495 | 1.68842 | 0.05289 |

\*Dwell-time distributions were fit with a single-exponential decay function, $y = y_0 + A_1 \exp(-t/\tau)$ . $\tau$ is the fitted decay time constant, and Fit SE is the standard error of the fitted $\tau$ estimate.
The number of dwell events is the total number of events included in the fit for each condition.

**Table S2.** Apparent first-binding kinetics of Polθ-H under the indicated RPA and DNA-substrate conditions.

| Condition | Amp (%)<br>[95% CI] | $k_{app}$ (s<sup>-1</sup>)<br>[95% CI] | Bootstrap SE (s<sup>-1</sup>) | N |
| --- | --- | --- | --- | --- |
| -RPA/<br>Polθ-H<sup>WT</sup> | 36.4<br>[30.8, 45.7] | 0.00468<br>[0.00312, 0.00630] | 0.00083 | 911 |
| +RPA<sup>WT</sup>/<br>Polθ-H<sup>WT</sup> | 83.2<br>[80.7, 85.6] | 0.01558<br>[0.01424, 0.01707] | 0.00074 | 963 |
| +RPA<sup>D-4A</sup>/<br>Polθ-H<sup>WT</sup> | 33.5<br>[12.1, 100.0] | 0.00103<br>[0.00026, 0.00389] | 0.00110 | 767 |
| +RPA<sup>C-3A</sup>/<br>Polθ-H<sup>WT</sup> | 35.9<br>[19.8, 100.0] | 0.00156<br>[0.00045, 0.00356] | 0.00089 | 1,118 |
| -RPA/<br>Polθ-H<sup>E691R</sup> | 36.5<br>[31.6, 44.3] | 0.00517<br>[0.00367, 0.00668] | 0.00075 | 915 |
| +RPA<sup>WT</sup>/<br>Polθ-H<sup>E691R</sup> | 35.7<br>[27.3, 57.6] | 0.00314<br>[0.00162, 0.00477] | 0.00083 | 1,043 |
| -RPA/<br>Polθ-H<sup>WT</sup> (5'end) | 42.9<br>[33.3, 74.8] | 0.00338<br>[0.00151, 0.00526] | 0.00096 | 635 |
| +RPA<sup>WT</sup>/<br>Polθ-H<sup>WT</sup> (5'end) | 29.2<br>[20.3, 82.8] | 0.00304<br>[0.00077, 0.00544] | 0.00118 | 621 |

\*A is the fitted plateau amplitude (%) of the binding curve, and $k_{app}$ is the apparent first-binding rate constant (s<sup>-1</sup>).
Values are reported as fitted estimates with 95% bootstrap confidence intervals; bootstrap SEs for $k_{app}$ are listed separately. N denotes the total number of DNA molecules analyzed for each condition. Bootstrap SEs and 95% confidence intervals were estimated from 2,000 bootstrap resamples, with molecules resampled within each independent experiment. Bootstrap SEs reflect within-experiment bootstrap uncertainty and do not represent SEM across independent experiments.

**Table S3.** Cryo-EM data collection statistics.

|  |  |
| --- | --- |
| Microscope | Thermo Fisher Titan Krios |
| Voltage (keV) | 300 |
| Camera | Falcon 4i |
| Magnification | 105,000 |
| Pixel size at detector (Å per pixel) | 0.94 |
| Defocus range (µm) | −1 to −1.8 |
| Energy filter slit width (eV) | 20 |
| Total electron exposure (e<sup>−</sup> Å<sup>−2</sup>) | 50.94 |
| Exposure rate (e<sup>−</sup> Å<sup>−2</sup> s<sup>−1</sup>) | 9.07 |
| Number of frames collected during exposure | 48 |
| Automation software | EPU |
| Micrographs collected (no.) | 9,741 |
| Micrographs used (no.) | 9,741 |
| Total extracted particles (no.) | 5,092,152 |
| **Polθ-H-RPA-ssDNA dimer (Map A)** |  |
| EMD-79697 |  |
| Final particles (no.) | 231,659 |
| Symmetry imposed | C1 |
| Resolution (global, Å) |  |
| FSC 0.143 masked | 3.4 |
| FSC 0.5 masked | 3.8 |
| Map sharpening *B* factor (Å<sup>2</sup>) | −110.6 |
| 3DFSC sphericity | 0.798 |
| Map sharpening methods | cryoSPARC |
| **Polθ-H (Map B)** |  |
| EMD-79704 |  |
| Final particles (no.) | 231,659 |
| Symmetry imposed | C1 |
| Resolution (global, Å) |  |
| FSC 0.143 masked | 3.2 |
| FSC 0.5 masked | 3.6 |
| Map sharpening *B* factor (Å<sup>2</sup>) | −104.5 |
| 3DFSC sphericity | − |
| Map sharpening methods | cryoSPARC, DeepEMhancer |
| **Polθ-H (Map C)** |  |
| EMD-79707 |  |
| Final particles (no.) | 231,659 |
| Symmetry imposed | C1 |
| Resolution (global, Å) |  |
| FSC 0.143 masked | 3.2 |
| FSC 0.5 masked | 3.8 |
| Map sharpening *B* factor (Å<sup>2</sup>) | −104.9 |
| 3DFSC sphericity | − |
| Map sharpening methods | cryoSPARC, DeepEMhancer |
| **RPA (Map D)** |  |
| EMD- 79716 |  |
| Final particles (no.) | 231,659 |
| Symmetry imposed | C1 |
| Resolution (global, Å) |  |
| FSC 0.143 masked | 3.4 |
| FSC 0.5 masked | 4.0 |
| Map sharpening *B* factor (Å<sup>2</sup>) | −113.9 |
| 3DFSC sphericity | - |
| Map sharpening methods | cryoSPARC, DeepEMhancer |
| **RPA (Map E)** |  |
| EMD- 79717 |  |
| Final particles (no.) | 231,659 |
| Symmetry imposed | C1 |
| Resolution (global, Å) |  |
| FSC 0.143 masked | 3.4 |
| FSC 0.5 masked | 4.0 |
| Map sharpening *B* factor (Å<sup>2</sup>) | -129.6 |
| 3DFSC sphericity | - |
| Map sharpening methods | cryoSPARC, DeepEMhancer |

**Table S4.** Model composition, refinement, and validation.

| Polθ-H-RPA-ssDNA dimer |  |
| --- | --- |
| PDB: 39RH |  |
| EMD- 79718 |  |
| **Model composition** |  |
| Non-hydrogen atoms (no.) | 39,220 |
| Protein residues (no.) | 2,407 |
| Ligands (no.) | 0 |
| Nucleotide residues (no.) | 25 |
| **Model Refinement** |  |
| Initial models used (PDB #) | 9BH8, 8RK2 |
| Refinement packages | Coot, PHENIX, ISOLDE |
| Model-Map scores |  |
| Cross-correlation coefficient | 0.72 |
| Model resolution (Å) | 3.5 |
| FSC threshold | 0.5 |
| Mean *B* factors (Å<sup>2</sup>) |  |
| Protein residues (no.) | 81.75 |
| Ligands (no.) | n/a |
| Nucleotide residues (no.) | 100.10 |
| R.m.s. deviations from ideal values |  |
| Bond lengths (Å) | 0.003 |
| Bond angles (°) | 0.567 |
| **Validation** |  |
| MolProbity score | 1.75 |
| CaBLAM outliers (%) | 1.29 |
| Clashscore | 8.09 |
| Poor rotamers (%) | 0.67 |
| C-beta deviations (%) | 0.00 |
| Ramachandran plot |  |
| Favored (%) | 95.53 |
| Allowed (%) | 4.43 |
| Outliers (%) | 0.04 |

